# A systematic cell size screen uncovers coupling of growth to division by the p38/HOG network in *Candida albicans*

**DOI:** 10.1101/094144

**Authors:** Adnane Sellam, Julien Chaillot, Jaideep Mallick, Faiza Tebbji, Julien Richard Albert, Michael A. Cook, Mike Tyers

## Abstract

Cell size is a complex trait that responds to developmental and environmental cues. Quantitative analysis of the size phenome in the pathogenic yeast *Candida albicans* uncovered 195 genes that markedly altered cell size, few of which overlapped with known size genes in other yeast species. A potent size regulator specific to *C. albicans* was the conserved p38/HOG MAPK module that mediates the osmotic stress response. Basal HOG activity inhibited the SBF G1/S transcription factor complex in a stress-independent fashion to delay the G1/S transition. The HOG network also governed ribosome biogenesis through the master transcriptional regulator Sfp1. Hog1 bound to the promoters and cognate transcription factors for both the G1/S and ribosome biogenesis regulons and thereby directly linked cell growth and division. These results illuminate the evolutionary plasticity of size control and identify the HOG module as a nexus of cell cycle and growth regulation.

## Introduction

A central and longstanding problem in cell biology is how cells maintain a uniform cell size, whether in single celled organisms or in the multitude of tissues of metazoans (1, 2). In most eukaryotes, attainment of a critical cell size appears to be necessary for commitment to cell division in late G1 phase, called Start in yeast and the Restriction Point in metazoans. This critical cell size threshold coordinates cell growth with cell division to establish a homeostatic cell size (1). The dynamic control of cell size facilitates adaptation to changing environmental conditions in microorganisms and therefore is essential to maximize fitness (3, 4). In the budding yeast *Saccharomyces cerevisiae,* the size threshold is dynamically modulated by nutrients. Pre-Start G1 phase cells grown in the optimal carbon source glucose pass Start at a smaller size if shifted to glycerol, whereas cells shifted from a poor to rich nutrient source pass Start at a larger size (1, 5). Nutrient conditions similarly dictate cell size in the fission yeast *Schizosaccharomyces pombe,* although control is primarily exerted at the G2/M transition (5). In metazoans, size control is important for tissue-specific functions and organ or organism size (6). Cell size is often perturbed in human disease, for example in diabetes, tuberous sclerosis, mitochondrial disorders, aneuploid syndromes, cancer and aging (1, 7). Notably, a loss of cell size homeostasis, termed pleomorphism, correlates with poor cancer prognosis (8).

Cell size is fundamentally dictated by the balance between cell growth and division. The analysis of small-sized mutants in yeast led to key insights into the cell cycle machinery (9-13). In all eukaryotes, cell division is controlled by the cyclin dependent kinases (CDKs), which serve to coordinate the replication and segregation of the genome (14). In *S. cerevisiae,* the G1 cyclins Cln1, Cln2 and Cln3 trigger Start, whereas the B-type cyclins Clb1-Clb6 catalyze replication and mitosis, all via activation of the same Cdc28 kinase catalytic subunit. The expression of ~200 genes at the end of G1 phase, most vitally *CLN1*/2, is controlled by transcription factor complexes composed of Swi4 and Swi6 (SBF), and Mbp1 and Swi6 (MBF). Activation of SBF/MBF depends primarily on the Cln3-Cdc28 kinase, the key target of which is Whi5, an inhibitor of SBF/MBF-dependent transcription (15, 16). Another transcriptional inhibitor called Nrm1 specifically inhibits MBF after Start but does not cause a marked size phenotype under conditions of nutrient sufficiency (16). Size control in *S. pombe* is exerted through inhibition of G2/M phase CDK activity by the Wee1 kinase, which is encoded by the first size control gene discovered (9, 11). Size is also partially regulated at Start in *S. pombe* through an SBF/MBF- like G1/S transcription factor complex and the Nrm1 inhibitor (17). The CDK-dependent control of G1/S transcription in metazoans is analogously mediated by the cyclin D-Rb-E2F axis (15, 18, 19).

Cell growth depends on the coordinated synthesis of protein, RNA, DNA and other macromolecules (1, 20, 21). The production of ribosomes consumes a large fraction of cellular resources and depends on an elaborate ribosome biogenesis machinery (1) that is controlled in part by the conserved TOR (<u>T</u>arget <u>O</u>f <u>R</u>apamycin) nutrient sensing network (6). Systematic size analysis in yeast uncovered many ribosome biogenesis (*Ribi*) genes as small size mutants, and revealed two master regulators of *Ribi* gene expression, the transcription factor Sfp1 and the AGC kinase Sch9, as the smallest mutants (22). These observations lead to the hypothesis that the rate of ribosome biogenesis is one metric that dictates cell size (23). Sfp1 and Sch9 are critical effectors of the TOR pathway and form part of a dynamic, nutrient-responsive network that controls the expression of *Ribi* and ribosomal protein *(RP)* genes (23). Sfp1 activity is controlled through its TOR-dependent nuclear localization (22-25) and is physically linked to the secretory system by its interaction with the Rab escort factor Mrs6 (24, 26). Sch9 is phosphorylated and activated by TOR, and in turn inactivates a cohort of repressors of *RP* genes called Dot6, Tod6 and Stb3 (27). The TOR network also controls size in *S. pombe* and metazoans (28).

Systematic genetic analysis has uncovered hundreds of genes that directly or indirectly affect cell size. Direct size analysis of all strains in the *S. cerevisiae* gene deletion collection uncovered a number of potent size regulators, including Whi5, Sfp1 and Sch9 (22, 29, 30), and revealed inputs into size control from ribosome biogenesis, mitochondrial function and the secretory system (23-26). Subsequent analyses of many of these size mutants at a single cell level have suggested that the critical cell size at Start may depend on growth rate in G1 phase and/or on cell size at birth (31, 32). Visual screens of *S. pombe* haploid and heterozygous deletion collections for size phenotypes also revealed dozens of novel size regulators, many of which altered size in a genetically additive fashion (33, 34). A large number of genes appear to influence size in metazoan species. A large-scale RNAi screen in *Drosophila melanogaster* tissue culture cells revealed hundreds of genes as candidate size regulators, including known cell cycle regulatory proteins (35). Despite overall conservation of the central processes that control cell growth and division, many functionally equivalent size regulators appear not to be conserved at the sequence level. For example, the G1/S transcriptional regulators SBF/MBF and Whi5 bear no similarly to the metazoan counterparts E2F and Rb, respectively (1). A number of TOR effectors are also poorly conserved at the sequence level, including the ribosome biogenesis transcription factors Sfp1 in yeast and Myc in metazoans (1).

*Candida albicans* is a diploid ascomycete yeast that is a prevalent commensal and opportunistic pathogen in humans. *C. albicans* is a component of the normal human flora, colonizing primarily mucosal surfaces, gastrointestinal and genitourinary tracts, and skin (36). Although most *C. albicans* infections entail non-life-threatening colonization of surface mucosal membranes, immunosuppressed patients can fall prey to serious infections, such as oropharyngeal candidiasis in HIV patients and newborns, and lethal systemic infections known as candidemia (37). Interest in *C. albicans* is not limited to understanding its function as a disease-causing organism, as it has an ecological niche that is obviously distinct from the classic model ascomycete *S. cerevisiae.* In this regard, *C. albicans* has served as an important evolutionary milepost with which to assess conservation of biological mechanisms, and recent evidence suggests a surprising extent of rewiring of central signalling and transcriptional networks as compared to *S. cerevisiae* (38-42).

In order to probe the conservation of the extensive size homeostasis network, we performed a quantitative genome-wide analysis of a systematic collection of gene deletion strains in *C. albicans.* Our results revealed an unexpected high degree of divergence between genes that affect size in *C. albicans* versus *S. cerevisiae* and uncovered previously undocumented regulatory circuits that govern critical cell size at Start in *C. albicans*. In particular, we delineate a novel stress-independent function of the p38/HOG MAPK network in coupling cell growth to cell division. Our genetic and biochemical analysis suggests that the HOG module directly interacts with central components of both the cell growth and cell division machineries in *C. albicans.* This systematic analysis thus provides insights into the general architecture and evolvability of the eukaryotic cell size control machinery.

## Results

### Analysis of the cell size phenome in *C. albicans*

The diploid asexual lifestyle of *C. albicans* complicates the discovery of loss-of-function phenotypes because both alleles must be inactivated (43). To uncover genes required for cell size homeostasis in *C. albicans,* we therefore directly screened four systematic collections of homozygous diploid gene deletion strains that encompassed 426 transcriptional regulators (41, 44), 81 kinases (40) and a general collection of 666 functionally diverse genes (45). We also screened a large set of 2360 strains from the GRACE (Gene Replacement and Conditional Expression) collection, which bear a gene deletion at one locus and an integrated tetracycline-regulated allele at the other locus (46). In total, 2630 viable mutant strains (2478 unique mutants) representing ~ 40 % of all predicted genes in *C. albicans* were individually assessed for their size distribution under conditions of exponential growth in rich medium. We additionally examined selected deletion strains of *C. albicans* orthologs of known size genes in *S. cerevisiae (sch9, pop2, ccr4, nrm1* and *CLN3/cln3)* that were not present in extant deletion collections **(Supplementary file 1)**. Clustering of size distributions across the cumulative datasets revealed distinct subsets of both large and small mutants, relative to the majority of mutants that exhibited size distributions comparable to those of wild-type (wt) control strains (**Figure 1A-D**). Mean, median and mode cell size were estimated for each mutant strain, and mutants were classified as large or small on the basis of a stringent cut-off of a 20 % increase or decrease in size as compared to the parental strain background. This cut-off value was determined based on a benchmark set of conserved small (*sch9*, *sfp1*) and large (*swi4*, *pop2*, *ccr4*) sized mutants for which size was reduced or increased at least 20 % as compared to parental strains. Based on this criterion, we identified 195 mutants that exhibited a size defect compared to their parental strain, comprised of 104 small sized *(whi)* and 91 large sized *(lge)* mutants **(Supplementary file 2 and 3)**.

**Figure 1.**
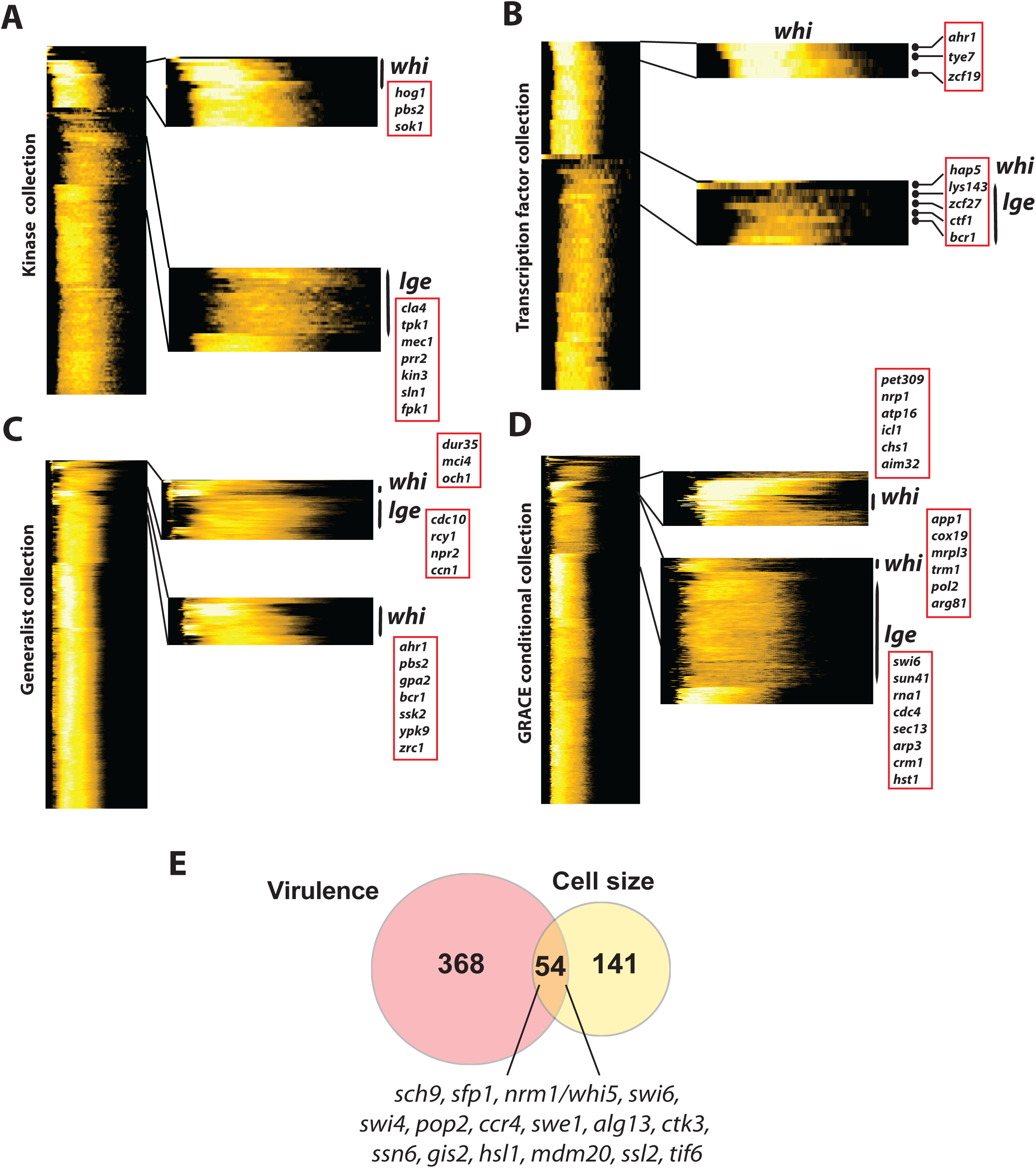
The cell size phenome of *C. albicans.* Clustergrams of size profiles of four different systematic mutant collections of *C. albicans:* **(A)** a set of 81 kinases (40); **(B)** a set of 166 transcription factors (41); **(C)** a general collection of 666 diverse genes (45); **(D)** the GRACE conditional collection of 2360 genes (46). Size distributions of each mutant (rows, represented as a heatmap) were normalized as percentage of total counts, smoothed by averaging over a seven-bin sliding window, and hierarchically clustered using a custom R script. **(E)** Overlap between *C. albicans* size and virulence phenotypes. Avirulent mutant phenotypes were obtained from CGD based on decreased competitive fitness in mice and/or reduced invasion and damage to host cells.

Gene Ontology (GO) enrichment analysis revealed that size mutants were predominantly defective in functions related to signalling, metabolism, transcriptional regulation, growth and cell cycle control (**Figure 1–figure supplement 1A**). For example, disruption of the HOG MAPK pathway (*hog1, pbs2, ssk2*), regulators of the G2/M phase transition *(swe1, gin4, hsl1, dbp11),* and protein localization to cell cortex *(mlc1, iqg1, gin4)* resulted in a small cell size phenotype. Conversely, mutants defective in functions related to the G1/S phase transition *(cln3, swi4, grr1)* and cytokinesis (*kin2*, *kin3*, *sec3*, *eng1*, *sun41*) caused a large cell size phenotype. As in *S. cerevisiae*, disruption of the central SBF (Swi4-Swi6) G1/S transcription factor complex increased cell size (**Figure 1–figure supplement 2E**), whereas mutation of the ribosome biogenesis regulators Sch9 and Sfp1 reduced cell size, as did inactivation of Cbf1, the major transcriptional regulator of ribosomal protein genes in *C. albicans* and other ascomycetes (47, 48).

Disruption of many other genes also perturbed size homeostasis in *C. albicans,* including genes implicated in cell wall structure and integrity *(gsc1*, *chs1, crh12, sun41, kre1, cbk1*), amino acid biosynthesis *(ssy1, car2, aro80, leu3, stp1, aco1),* and cellular respiration *(nuo1, mci4, cox19, hap2, hap43),* amongst many other processes **(Supplementary file 3)**. Interestingly, 57 of the 195 size mutants identified by our screen have been shown previously to be required for pathogenesis (p-value=1.23e-10). This set of genes included those with functions in transcriptional control of biofilm and invasive filament formation (*ndt80*, *efg1, nrg1, ace2, zcf27)* as well as known adhesion genes *(ahr1, efg1).* This overlap suggested that cell size homeostasis may be elemental to *C. albicans* fitness inside the host (**Figure 1E**, **Figure 1–figure supplement 1A** and **Supplementary file 7**).

### Limited overlap of the fungal cell size phenome across species

*C. albicans* and *S. cerevisiae* represent yeast genera separated by ~70M years of evolution (49) and share the morphological trait of budding, as well as core cell cycle and growth regulatory mechanisms (50, 51). However, recent evidence has uncovered an extensive degree of rewiring of both cis-transcriptional regulatory circuits and signalling pathways across many cellular and metabolic processes between the two yeasts (38, 40, 48, 52). To assess the extent of conservation and plasticity of the size phenome between the two species, genes that affected cell size in *C. albicans* were compared to their corresponding orthologs in *S. cerevisiae.* Surprisingly, we found minimal overlap between homozygous size mutants in both species (**Figure 2A-B**). Only four small size mutants we tested were common between *C. albicans* and *S. cerevisiae,* namely the master ribosome biogenesis regulators Sfp1 and Sch9, the CDK-inhibitory kinase Swe1 and the G1/S transcriptional repressor Nrm1. Similarly, only four large size mutants were shared, namely the SBF subunits Swi4 and Swi6 and two components of the CCR-NOT transcriptional complex, Ccr4 and Pop2 (**Figure 2B**). Heterozygous loss of the G1 cyclin Cln3 also caused a large size phenotype (53), but this gene was not included in the deletion collections we screened. It is likely that a number of other size regulators not represented in our collections may also be shared between *C. albicans* and *S. cerevisiae*, but nonetheless the limited degree of overlap in size regulators was strikingly limited considering that our collections spanned ~40 % of all *C. albicans* genes. Unexpectedly, six *C. albicans* small sized mutants *(gis2, ctk3, ssn6, hsl1, mdm20, sst2)* exhibited a large phenotype in *S. cerevisiae,* and conversely two *C. albicans* large mutants *(alg13* and *tif6)* caused a small cell size in *S. cerevisiae.* To confirm these disparities in the size control network independently of the particular cut-offs used to define large and small mutants in different studies, we examined the relationship between mode size of all screened *C. albicans* mutants and their counterparts in *S. cerevisiae.* Overall, no correlation was observed between the size phenomes of the two species across any of the *C. albicans* deletion collections (**Figure 2C-G**).

**Figure 2.**
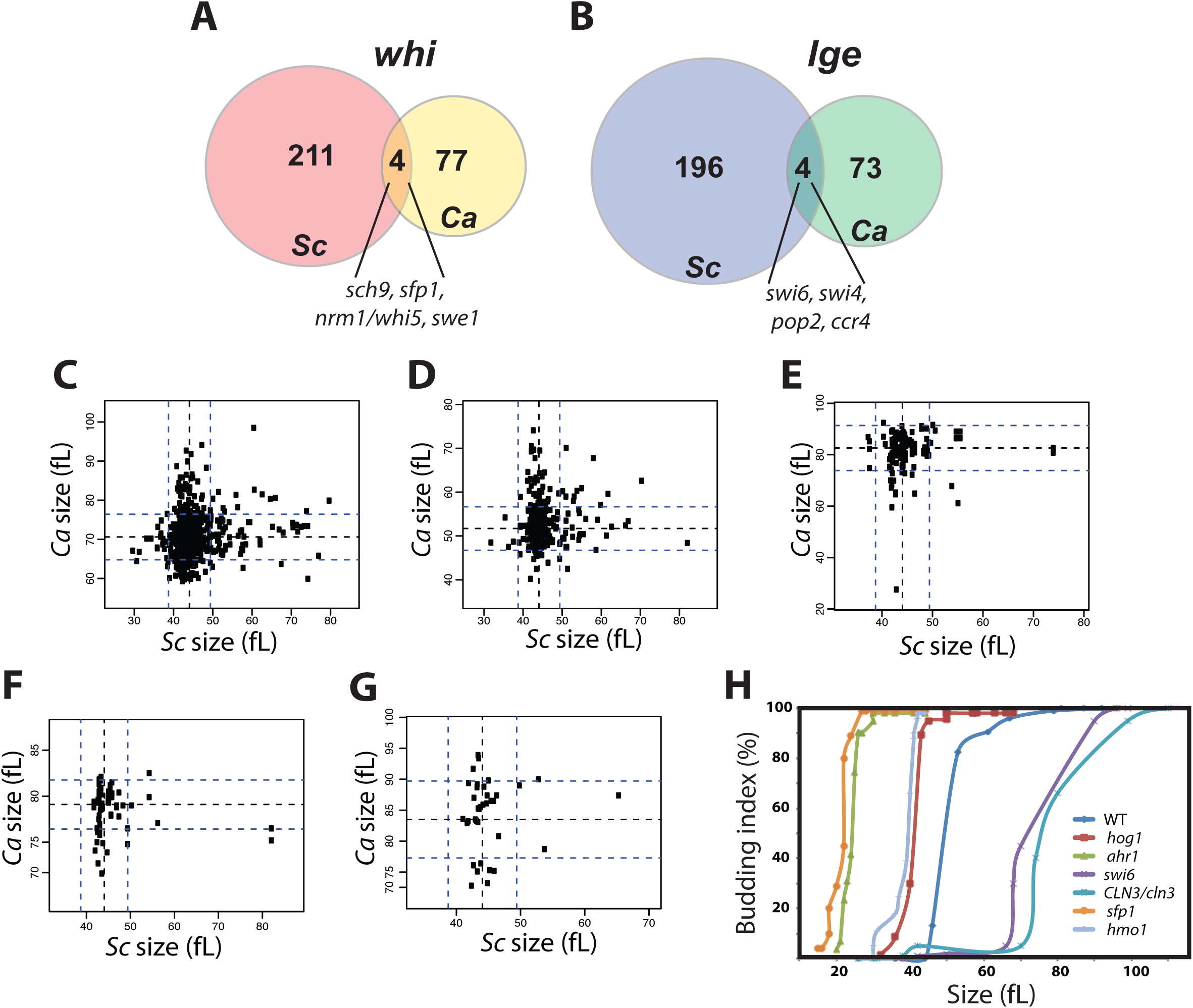
Comparative analysis of two systematic fungal cell size phenomes. Overlap between small **(A)** and large sizes strains **(B)** for *C. albicans* (Ca; this study) and *S. cerevisiae* (Sc) (22). **(C-G)** Scatter plots of mean sizes of *S. cerevisiae* deletion strains versus their counterparts in *C. albicans* in different mutant collections: **(C)** GRACE (46); **(D)** general (45); **(E)** kinases (40); **(F)** transcriptional regulators (44); **(G)** transcription factors (41). Dashed lines indicate standard deviation. **(H)** Size regulators in *C. albicans* act at Start. Early G1-phase cells of different potent size mutants and wt (SN250 (45)) were isolated by centrifugal elutriation, released into fresh YPD medium and monitored for bud emergence and cell size at 10 min intervals until the entire population was composed of budded cells.

We also compared size genes in *C. albicans* to those previously identified in a systematic visual screen for size mutants in the *S. pombe* haploid deletion strain collection (33). For the 18 validated size regulators in this *S. pombe* screen, a total of 10 had obvious *C. albicans* orthologs in our datasets. Comparison of size phenotypes for these 10 genes revealed that only the CDK-inhibitory kinase *swe1* (*wee1* in *S. pombe)* caused an analogous small size phenotype in both species. Overall, these comparative analyses demonstrated that relatively few specific gene functions required for cell size homeostasis were conserved between *C. albicans, S. pombe* and *S. cerevisiae.*

### Novel Start regulators in *C. albicans*

Previous work has shown that disruption of cell growth rate is often accompanied by a small cell size phenotype, for example by mutations in *RP* or *Ribi* genes (22, 54). To identify *bona fide* negative Start regulators, as opposed to mere growth rate-associated effects, doubling times were determined for the 104 homozygous small size mutants identified in our screens **(Supplementary file 3)**. Mutants that exhibited a greater than 10 % increase in doubling time as compared to the wt controls were enriched in functions associated with macromolecular synthesis and were removed from subsequent consideration for this study. As expected, amongst the 63 remaining candidates predicted to more directly couple growth to division **(Supplementary file 4)**, we recovered three known conserved repressors of Start, namely Sfp1, Sch9 and Swe1. Strikingly, the candidate set of Start regulators in *C. albicans* contained many conserved genes that do not affect size in *S. cerevisiae,* including transcription factors (Hmo1, Dot6), the kinase Sok1, and components of the HOG MAPK pathway (Ssk2, Pbs2 and Hog1). Other novel potent candidate regulators of Start that were unique to *C. albicans* included genes with functions related to nitrogen metabolism (Aro80, Dur35), fatty acid biosynthesis (Ino4, Asg1) and iron metabolism (Hap2, Hap43). We also observed that loss of the F-box protein Grr1 resulted in a small size, consistent with the fact that this SCF ubiquitin ligase subunit eliminates the G1 cyclin Cln3 in *C. albicans* (55). While Grr1-mediated elimination of Cln3 is conserved in *S. cerevisiae,* the *grr1* deletion does not have a small size phenotype, in part because Cln3 is redundantly targeted by the F-box protein Cdc4 in this yeast (56).

We demonstrated the effect of six *C. albicans* size regulators on the timing of Start by assessing the correlation between size and bud emergence in a synchronous early G1 phase population of cells obtained by centrifugal elutriation. We used this assay to determine the effect of three potent novel size control mutants that conferred a small size phenotype *(ahr1*, *hog1*, *hmo1*) and, as a control, disruption of a conserved known regulator of Start *(sfp1).* Additionally, the Start transition in *C. albicans* was characterized for two large size mutants *(swi6* and *cln3).* The critical cell size of the four small sized mutants *ahr1, hog1, hmo1* and *sfp1* was markedly reduced as compared to the wt parental strain (**Figure 2H**). Conversely, as expected Start was delayed in the large sized mutants *CLN3/cln3* and *swi6.* These results demonstrate that the transcription factors Ahr1 and Hmo1, and the MAPK Hog1 are novel *bona fide* repressors of Start in *C. albicans,* and that aspects of the Start machinery appear to have diverged between *C. albicans* and *S. cerevisiae.*

### Basal activity of the HOG MAPK pathway delays Start

A *hog1* mutant strain had a median size that was 30 % smaller than its congenic parental strain, at 50.2 and 71.7 fL respectively (Figure 3A and 3C). To ascertain that this effect was mediated at Start, we evaluated two hallmarks of Start, namely bud emergence and the onset of SBF-dependent transcription as a function of cell size in synchronous G1 phase cells obtained by elutriation. As assessed by mode size of cultures for which 25 % of cells had a visible bud, the *hog1* mutant passed Start after growth to 41 fL, whereas a parental wt control culture passed Start at a much larger size of 55 fL (**Figure 3F**). Importantly, in the same experiment, the onset of G1/S transcription was accelerated in the *hog1* strain as judged by the peak in expression of the two representative G1 transcripts, *RNR1* and *PCL2* (**Figure 3G-H**). These results demonstrated that the Hog1 kinase normally acts to delay the onset of Start.

**Figure 3.**
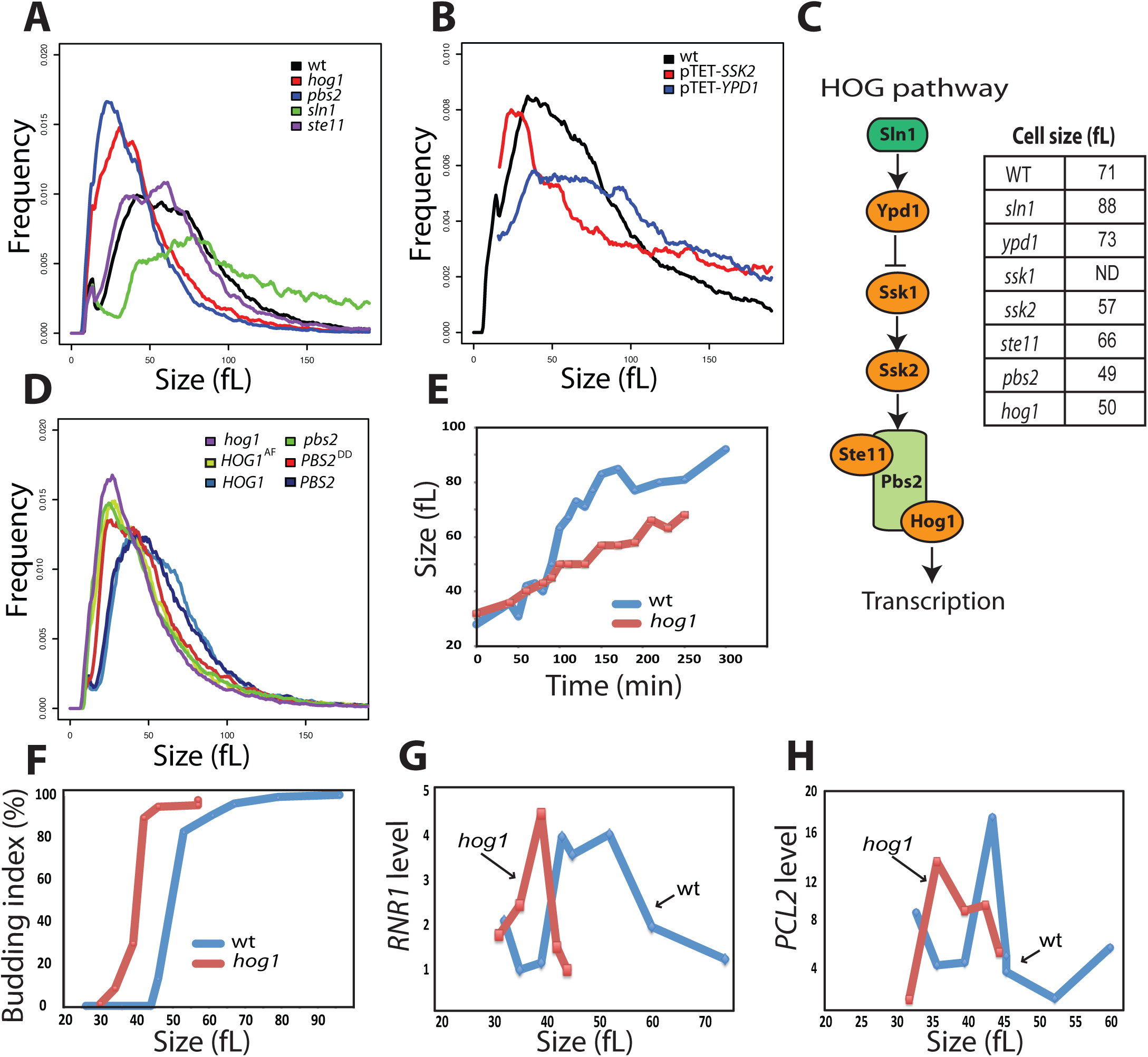
Basal activity of HOG pathway is required for normal Start onset and cell size homeostasis. **(A-B)** Size distributions of different mutant strains for the HOG pathway in *C. albicans.* **(C)** Schematic of the canonical HOG pathway in *C. albicans* and summary of mode size for each mutant strain. The *ssk1* strain exhibited constitutive filamentation that precluded size determination (ND = not determined). **(D)** Mutation of the two activating phosphorylation sites on Hog1 (T174A and Y176F, termed AF) and Pbs2 (S355D and T359D, termed DD) confers a small size phenotype. **(E-H)** Acceleration of Start in a *hog1* strain. **(E)** Elutriated G1 phase daughter cells were released into fresh media and assessed for size as a function of time, **(F)** bud emergence as a function of size and **(G-H)** G1/S transcription. *RNR1* and *PCL2* transcript levels were assessed by quantitative real-time PCR and normalized to *ACT1* levels.

Other main elements of the HOG pathway, namely the MAPKK Pbs2, the MAPKKK Ssk2, the phospho-relay mediator Ypd1 and the two-component transducer Sln1, were also required for normal cell size homeostasis (**Figure 3A-C**). Disruption of the upstream negative regulators Ypd1 and Sln1 caused a large size whereas mutation of the core MAPK module (Ssk2, Pbs2 and Hog1) caused a small size phenotype. As the cultures for these experiments were grown in constant normo-osmotic conditions, we inferred that the effect of the HOG module on cell size was unrelated to its canonical role in the osmotic stress response. Consistent with this interpretation, mutation of the known osmotic stress effectors of the HOG pathway in *C. albicans*, namely the glycerol biosynthetic genes *GPD1, GPD2* and *RHR2* (57, 58), did not cause a cell size defect (**Figure 3–figure supplement 1** and **Supplementary file 1**). To address whether basal activity of the HOG MAPK module might be required for size control, we tested the effect of phosphorylation site mutants known to block signal transmission. Mutation of the activating phosphorylation sites on either Hog1 (Thr174 and Tyr176) or Pbs2 (Ser355-and Thr359) to non-phosphorylatable residues phenocopied the small size of *hog1* and *pbs2,* respectively (**Figure 3D**). This result demonstrated that a basal level of Hog1 and Pbs2 activity was required for Start repression under non-stress conditions. To examine the possible role of the HOG pathway in communicating nutrient status to the Start machinery, the effects of different carbon sources on cell size were assessed in *hog1* and wt strains. Cell size was reduced on poor carbon sources in the *hog1* strain to the same extent as the wt strain, suggesting that the HOG module was not required for carbon source regulation of cell size (**Figure 3–figure supplement 2**). These results demonstrate that the HOG module relays a stress- and carbon source-independent signal for size control to the Start machinery in *C. albicans.*

Previous genome-wide screens in *S. cerevisiae* failed to uncover a role for the HOG pathway in size control (22, 29-31). To confirm these results, cell size distributions of HOG pathway mutants in *S. cerevisiae (hog1, pbs2, ssk1, ssk2, opy2* and *sho1* strains) were assessed in rich medium. None of the *S. cerevisiae* mutants had any discernable size defect as compared to a parental wt strain (**Figure 3–figure supplement 3**).

### Hog1 acts upstream of the SBF transcription factor complex

Cln3-dependent activation of the Swi4-Swi6 transcriptional complex drives G1/S progression in both *S. cerevisiae* and *C. albicans* (51, 53, 59, 60) such that *CLN3/cln3, swi6* and *swi4* mutants all exhibited large size and a G1 phase delay (**Figure 1–figure supplement 2E**, **Figure 2H** and 4A). To examine the functional relationship between the HOG pathway and these canonical Start regulators, we characterized their genetic interactions by size epistasis. We observed that the small size of a *hog1* mutant strain was partially epistatic to the large size of the heterozygous *CLN3/cln3* mutant (**Figure 4A**), suggesting that the HOG pathway may function in parallel to Cln3. Deletion of *SWI4* in the *hog1* mutant strain resulted in large size comparable to that of *swi4* mutant, suggesting that Hog1 acts upstream of Swi4 to inhibit Start (**Figure 4B**). In support of this finding, co-immunoprecipitation assays revealed that Hog1 physically interacted with Swi4 in a rapamycin-sensitive manner and that the Hog1-Swi4 interaction was insensitive to osmotic stress (**Figure 4C**). In *C. albicans,* the Nrm1 inhibitor is known to interact with the SBF complex to repress the G1/S transition (61), and consistently a *nrm1* mutant exhibited a reduced cell size (**Figure 4D**). We found that a *nrm1 hog1* double mutant had a smaller size than either of the *nrm1* or *hog1* single mutants, suggesting that Nrm1 and Hog1 act in parallel pathways to inhibit G1/S transcription (**Figure 4D**). Collectively, these genetic and biochemical results identified Hog1 as a new regulator of SBF in *C. albicans,* and suggested that Hog1 may transmit signals from the TOR growth control network to the G1/S machinery.

**Figure 4.**
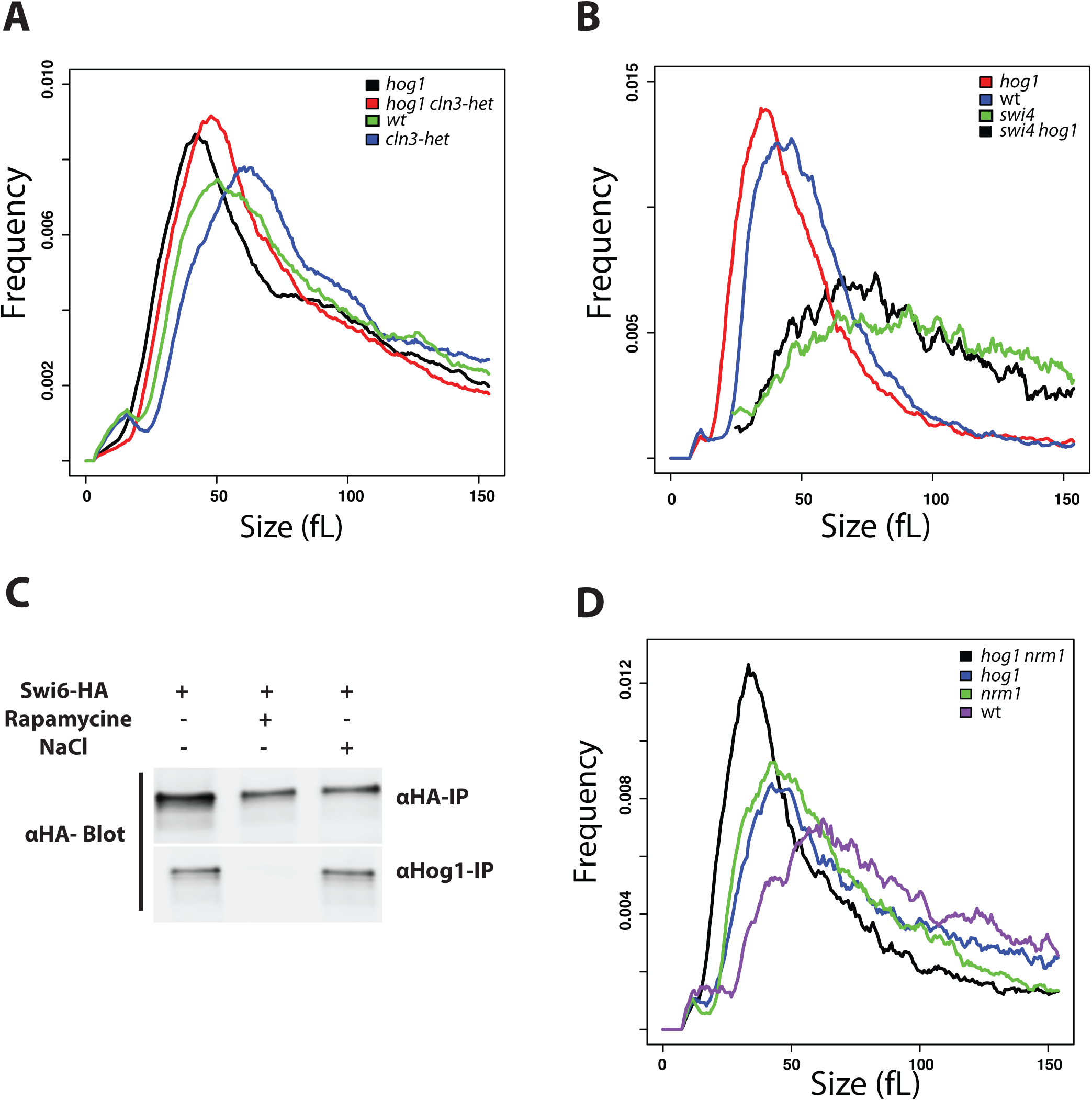
Genetic interactions between the HOG pathway and the G1/S transcriptional machinery. **(A)** Additive effect of *hog1* and *Cln3/cln3* mutations on cell size. The wt strain was the SN148-Arg+ parental background. **(B)** A *swi4* mutation is epistatic to a *hog1* mutation for cell size. The wt was the SN250. **(C)** Co-immunoprecipitation assays for Hog1 and Swi4. Cultures were treated as indicated with rapamycin at 0.5 μg/ml and NaCl at 0.5 M for 30 min. **(D)** Additive effect of *hog1* and *nrm1* mutations on cell size. The wt strain was the SN250.

### The Ptc1 and Ptc2 phosphatases control Start via basal Hog1 activity

MAPK activity is antagonized by the action of serine/threonine- (Ser/Thr) phosphatases, tyrosine (Tyr) phosphatases, and dual specificity phosphatases that are able to dephosphorylate both Ser/Thr and Tyr residues (62). In *S. cerevisiae,* after adaptation to osmotic stress, components of the HOG pathway are dephosphorylated by both protein Tyr phosphatases and type 2C Ser/Thr phosphatases (62, 63). In *C. albicans,* recent work has identified the two Tyr phosphatases Ptp2 and Ptp3 as modulators of the basal activity of Hog1 (64). A prediction of the HOG-dependent size control model is that disruption of the phosphatases that modulate Hog1 basal activity should cause a large cell size. However, none of the Tyr-phosphatase single mutants *ptp1, ptp2* or *ptp3,* nor a *ptp2 ptp3* double mutant exhibited a noticeable cell size defect (**Figure 5B** and **Supplementary file 1**). In contrast, deletion of the type 2C Ser/Thr phosphatase Ptc2 conferred a median size of 84.9 fL, which was 24 % larger than the parental wt control size of 68 fL, while a *ptc1 ptc2* double mutant strain had an even larger size of 90.5 fL (**Figure 5A**). To confirm that the large size phenotype of the *ptc* mutants was mediated directly via effects on Start, we evaluated the critical cell size of both *ptc2* and *ptc1 ptc2* mutants in elutriated G1 cells. Whereas wt control cells passed Start at 49 fL, the critical cell size of the *ptc2* and *ptc1 ptc2* mutant strains was increased by 59 % to 78 fL and 87 % to 92 fL, respectively (**Figure 5C**). To determine whether Hog1 is an effector of Ptc1 and Ptc2 at Start, we examined the epistatic relationship between the *hog1* and *ptc1 ptc2* mutations. The size of the *hog1 ptc1 ptc2* triple mutant was identical to that of *hog1* single mutant, indicating that Hog1 functions downstream of Ptc1 and Ptc2 for the control of cell size (**Figure 5D**). These data suggested that Ptc1 and Ptc2 phosphatases may modulate the basal phosphorylation state of Hog1 to govern the timing of Start onset and critical cell size.

**Figure 5.**
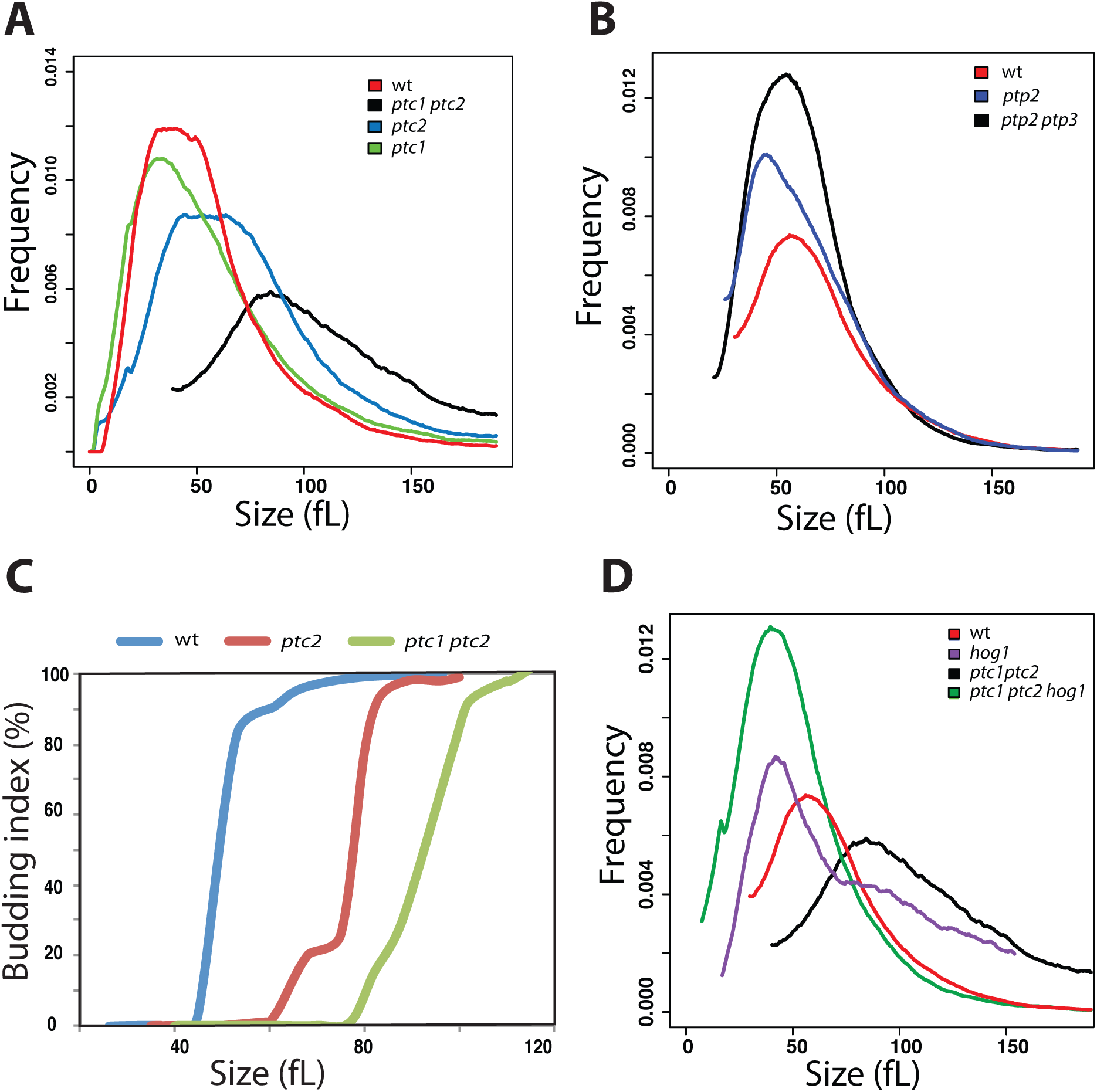
Ptc1 and Ptc2 control Start via Hog1. **(A)** Size distributions of a wt strain (SN250) and *ptc1, ptc2 and ptc1 ptc2* deletion mutants. **(B)** Size distributions of a wt strain (SN250), a *ptp2* single mutant, and a *ptp2 ptp3* double mutant. **(C)** Start is delayed in *ptc* mutants. Elutriated G1 phase daughter cells were released into fresh media and monitored for bud emergence as a function of size. **(D)** The small cell size of a *hog1* mutant is epistatic to the large size of a *ptc1 ptc2* double mutant.

### Hog1 activates ribosome biosynthetic gene transcription and inhibits G1/S transcription

To explore the role of Hog1 at Start, we assessed genome-wide transcriptional profiles using custom microarrays. G1 phase cells for *hog1* mutant and wt strains were collected by centrifugal elutriation, followed by microarray analysis of extracted total RNA. Gene set enrichment analysis of transcriptional profiles (65, 66) revealed that the *hog1* strain was defective in expression of genes that function in protein translation, including members of the 48S/43S translation initiation complex, structural components of the small and large subunits of the ribosome, and tRNA-charging components (**Figure 6A** and **Supplementary file 5**). Transcription of genes that function in mitochondrial transport, the tricarboxylic acid cycle, protein degradation by the 26S proteasome and respiration were also downregulated in a *hog1* deletion. Conversely, the G1/S transcriptional program (51) was hyperactivated in a *hog1* mutant, consistent with the above results for *RNR1* and *PCL2.* These results suggested that Hog1 activates multiple processes that underpin cellular growth in addition to its role as a negative regulator of the G1/S transcriptional program.

**Figure 6.**
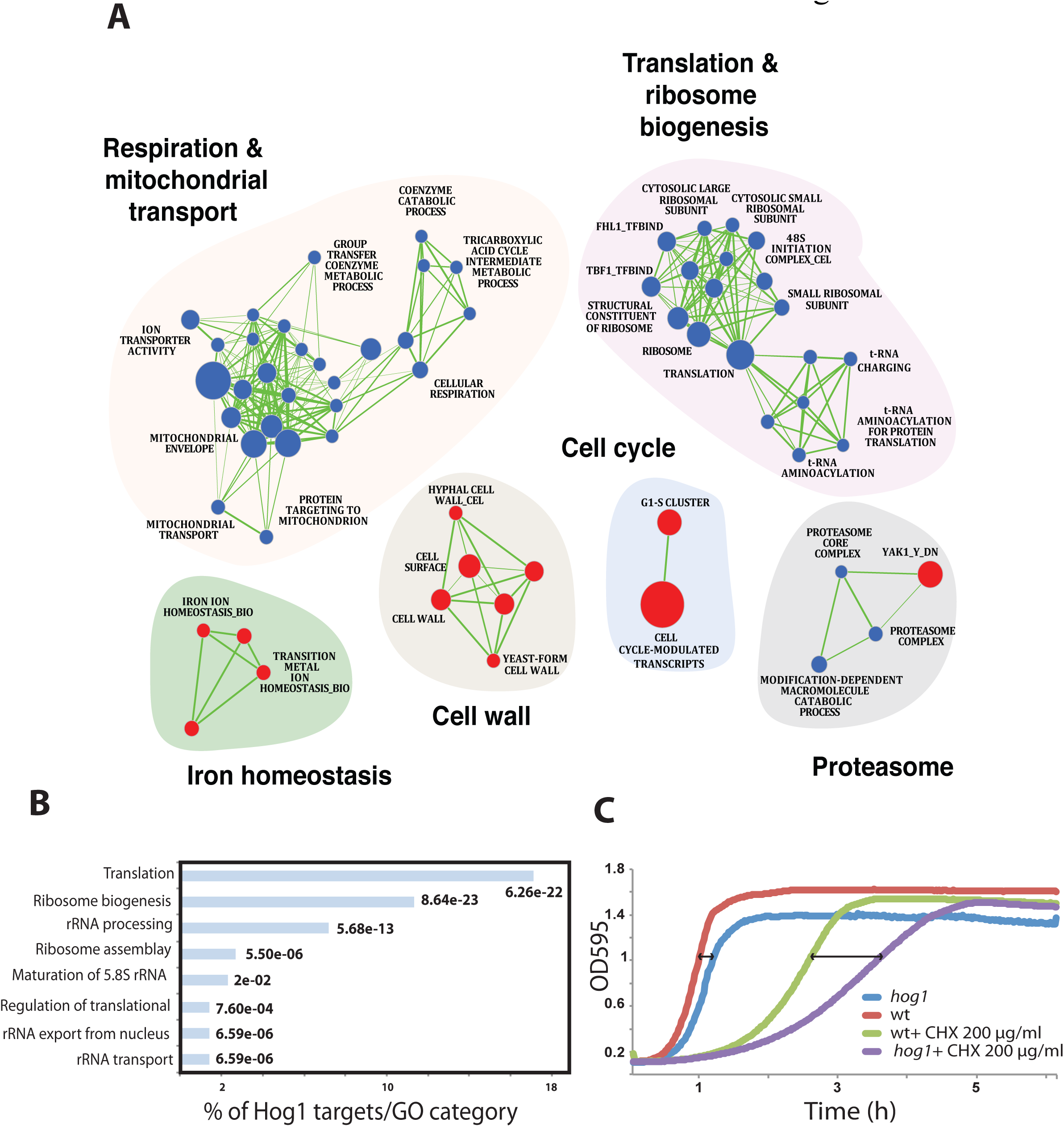
A Hog1-dependent transcriptional program in G1 phase cells. **(A)** GSEA analysis of differentially expressed genes in a *hog1* mutant relative to a congenic wt strain (SN250). Cells were synchronized in G1 phase by centrifugal elutriation and released in fresh YPD medium for 15 min and analyzed for gene expression profiles by DNA microarrays. Up-regulated (red circles) and down-regulated (blue circles) transcripts are shown for the indicated processes. The diameter of the circle reflects the number of modulated gene transcripts in each gene set. Known functional connections between related processes are indicated (green lines). Images were generated in Cytoscape with the Enrichment Map plug-in. **(B)** Genome-wide promoter occupancy of Hog1 in G1 phase cells. Gene categories bound by Hog1 were determined by GO term enrichment. *p*-values were calculated using hypergeometric distribution. **(C)** Growth rate and cycloheximide (CHX 200 μg/ml) sensitivity of wt and *hog1* mutant strains.

It has been previously reported that Hog1 in *S. cerevisiae* and its ortholog p38 in humans directly bind and activate downstream transcriptional target genes (67-72). In *S. cerevisiae,* Hog1 thus associates with DNA at stress-responsive genes and is required for recruitment of general transcription factors, chromatin modifying activities and RNA Pol II (68, 71, 73, 74). However, although mechanisms of Hog1-dependent transcription have been investigated under osmotic stress conditions, the function of this kinase in normal growth conditions in the absence of stress has not been explored. In order to assess whether Hog1 might directly regulate gene expression relevant to cell size control in *C. albicans,* we profiled the genome-wide localization of Hog1 in G1 phase cells obtained by centrifugal elutriation from TAP-tagged Hog1 and untagged control strains. Hog1 binding sites in the genome were determined in duplicate by chromatin immunoprecipitation and microarray analysis (ChIP-chip). These experiments revealed that Hog1^TAP^ was significantly enriched at 276 intergenic regions and 300 ORFs when compared to the untagged control **(Supplementary file 6**). The ORF and promoter targets of Hog1 were strongly represented for translation and *Ribi* genes (**Figure 6B**), in accord with the above expression profiles. These data suggested that Hog1 may directly activate expression of the *Ribi* regulon and other translation-associated genes. The strong enrichment for Hog1 at translation and *Ribi* loci suggested that Hog1 may be required for maximal translational capacity as G1 phase cells approach Start. Consistently, we observed that a *hog1* mutant exhibited increased sensitivity to the protein translation inhibitor cycloheximide as compared to a wt strain (**Figure 6C**). These results suggested that Hog1 may directly activate ribosome biogenesis and protein translation as cells approach Start.

### Hog1 is required for Sfp1-dependent gene expression and recruitment to target promoters

Based on the conserved role of the Sfp1 transcription factor and the kinase Sch9 in ribosome biogenesis and cell size control in *C. albicans*, we examined genetic interactions between these factors and the HOG pathway. To identify potential epistatic interactions, we overexpressed *SCH9* or *SFP1* in a *hog1* strain. The overexpression of *SFP1* but not *SCH9* restored the *hog1* strain to a near wt cell size distribution (**Figure 7A**). In contrast, overexpression of *HOG1* in either an *sfp1* or *sch9* had no effect on the small size of either strain (data not shown). These results suggested that Sfp1 might act downstream of Hog1. Consistent with this interpretation, we found that the gene expression defects of six *Ribi* and translation genes *(RPS12, RPS28B, RPS32, EIF4E* and *TIF6)* in a *hog1* strain were rescued by the overexpression of *SFP1* (**Figure 7B**).

**Figure 7.**
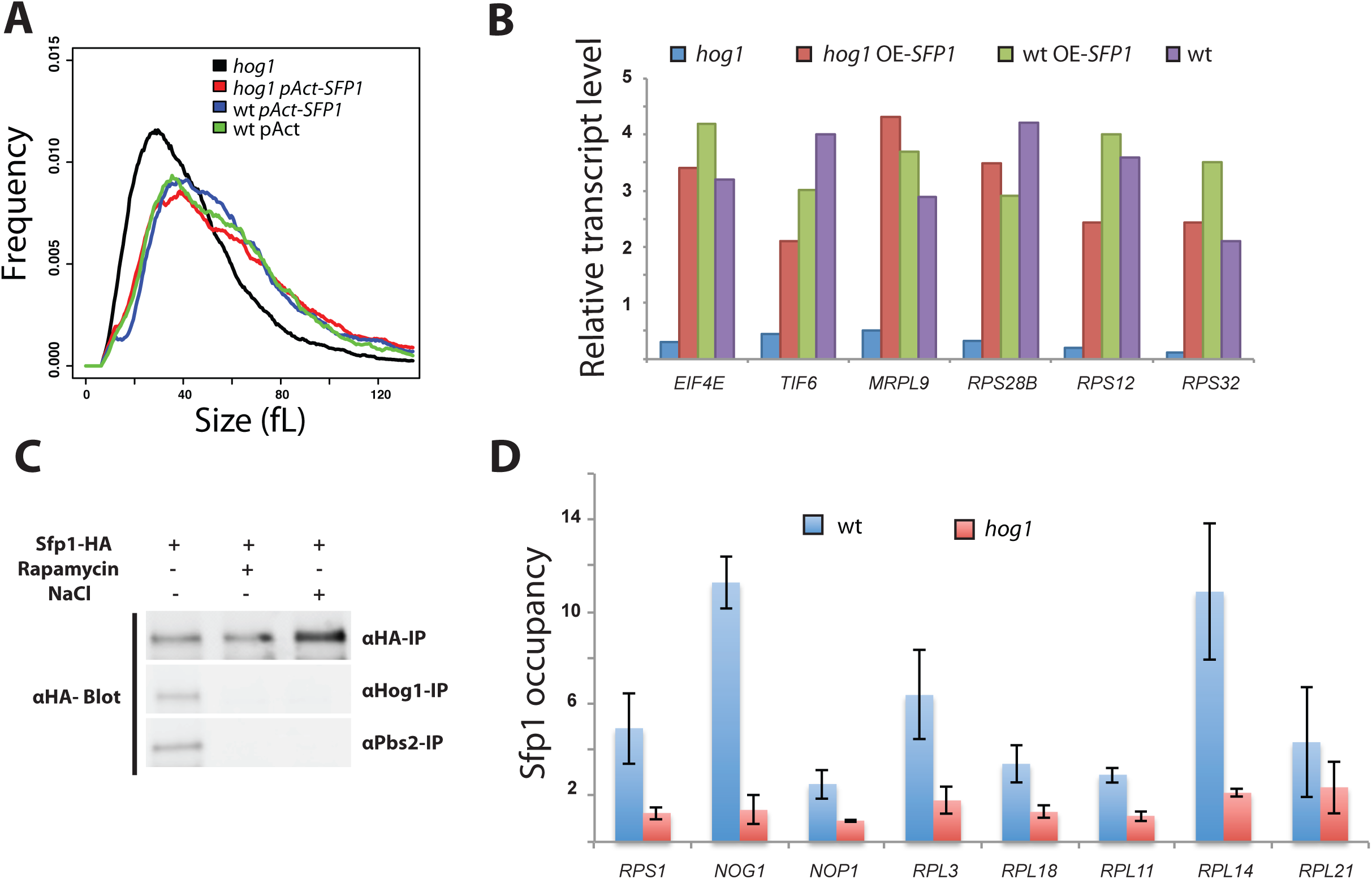
Hog1-dependent recruitment of Sfp1 to promoter DNA. **(A)** Size distributions of wt (wt-pAct), *hog1* (*hog1/hog1*), *sfp1*-overexpression (wt/pAct-sfp1) and *hog1 sfp1-*overexpression (hog1/hog1/pAct-sfp1) strains. **(B)** Increased *sfp1* dosage restores expression of representative *Ribi* and RP transcripts in a *hog1* mutant strain. Relative expression levels of the six transcripts were assessed by real-time qPCR as normalized to *ACT1.* Values are the mean from two independent experiments. **(C)** Sfp1 interactions with Pbs2 and Hog1. Anti-HA immunopreciptiates from a strain bearing an integrated SFP1^HA^ allele grown in the absence or presence of NaCl (0.5 M) or rapamycin (0.5 μg/ml) were probed with anti-HA, anti-Hog1 or anti-Pbs2 antibodies. **(D)** Reduced Sfp1 localization to *Ribi* gene promoters in a *hog1* mutant strain. Values are the mean from three independent ChIP-qPCR experiments for each indicated promoter.

Given the apparent genetic relationship between Hog1 and Sfp1, we examined whether the two proteins physically interacted. We evaluated the interaction at endogenous levels using a chromosomal HA-tagged Sfp1 allele and polyclonal antibodies that recognize Pbs2 and Hog1. Capture of Sfp1^HA^ from cell lysates followed by antibody detection revealed that Sfp1 interacted with both Pbs2 and Hog1 (**Figure 7C**). Notably, the Sfp1 interaction with both Hog1 and Pbs2 was abolished by either osmotic stress or rapamycin (**Figure 7C**). These results suggested that the timing of Start may be governed in part by modulation of the Hog1-Sfp1 interaction by stress and nutrient signals.

We then examined whether Sfp1 played an analogous role in Start control in *C. albicans* as in *S. cerevisiae.* As described above, an *sfp1* deletion strain was extremely small and passed Start at only 42 % of wt size (**Figure 2H** and **Figure 2–figure supplement 1A**). Consistently, transcriptional profiles of a strain bearing a tetracycline-regulated allele of *SFP1* demonstrated that expression of the *Ribi* regulon was partially Sfp1-dependent (**Figure 7–figure supplement 1A**). We also found that an *sfp1* strain was as sensitive to the protein translation inhibitor cycloheximide as a *hog1* strain (**Figure 7–figure supplement 1B**). These data demonstrated that Sfp1 is a transcriptional activator of *Ribi* genes and a negative regulator of Start in *C. albicans.*

The finding that both Hog1 and Sfp1 controlled the expression of *Ribi* genes, together with the finding that Hog1 acted upstream of Sfp1, led us to hypothesize that Hog1 might be required for the recruitment of Sfp1 to its target genes. To test this hypothesis, we used ChIP-qPCR to measure *in vivo* promoter occupancy of Sfp1^HA^ at eight representative *Ribi* and RP genes that were also bound by Hog1. While Sfp1 was detected at each of these promoters in a wt strain the ChIP signals were abrogated in the *hog1* mutant strain (**Figure 7D**). From these data, we concluded that Sfp1 regulates the *Ribi* regulon in a Hog1-dependent manner, and that the HOG module lies at the interface of the G1/S transcription and growth control machineries in *C. albicans*

## Discussion

This systematic genetic analysis of size control in *C. albicans* represents the first genome-wide characterization of the mechanisms underlying regulation of growth and division in a pathogenic fungus. As is the case for other species that have been examined to date, cell size in *C. albicans* is a complex trait that depends on diverse biological processes and many hundreds of genes (22, 29-31, 33, 35). Of particular note, our screen and subsequent molecular genetic analysis uncovered a novel function for the Hog1 as a critical nexus of the growth and division machineries. The HOG module thus represents a long-sought direct linkage between cell growth and division (**Figure 8**).

**Figure 8.**
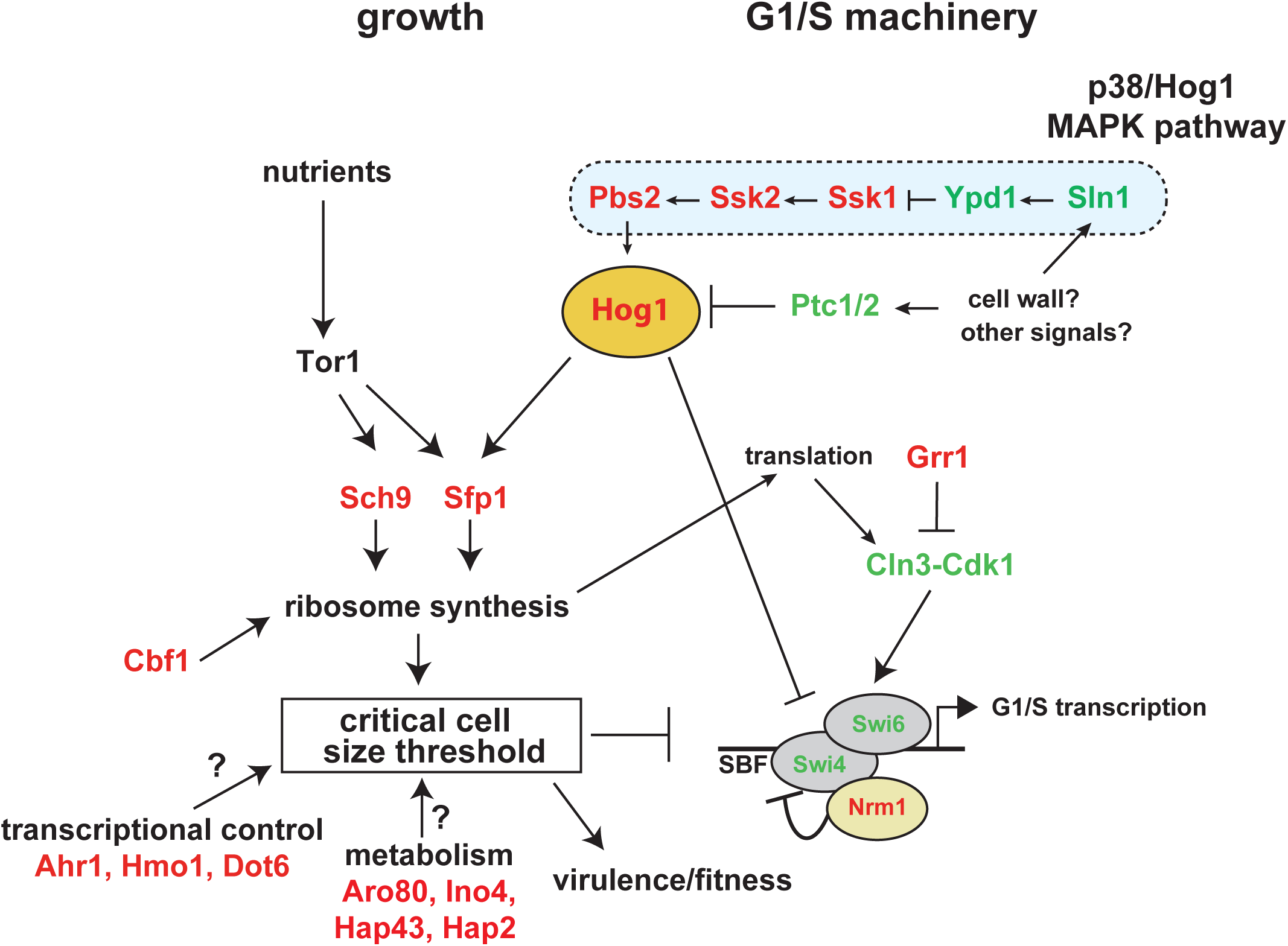
Architecture of the Start machinery in *C. albicans.* Hog1 inhibits the SBF G1/S transcription factor complex and in parallel controls Sfp1 occupancy of *Ribi* gene promoters, and thereby directly links growth and division. Basal activity of Hog1 is modulated by the phosphatases Ptc1 and Ptc2 to govern the timing of Start onset. Parallel Start pathways revealed by genetic interactions with Hog1, as well as other prominent size control genes in *C. albicans* revealed by size screens, are also indicated. Size regulators for which gene inactivation led to small and large size phenotypes are indicated in red and green, respectively.

### Conservation and divergence of cell size control mechanisms

Inactivation of many genes that control ribosome biogenesis and protein translation in *C. albicans* resulted in a small cell size, consistent with the notion that the rate of ribosome biogenesis is a component of the critical size threshold (1, 23). Mutation of the key conserved *Ribi* regulators Sch9 and Sfp1, as well as other *Ribi* factors, thus reduced cell size in *C. albicans.* Previous studies have shown that several RP and *Ribi* trans-regulatory factors have been evolutionarily rewired in *C. albicans* compared to *S. cerevisiae* (47). Consistently, we found that deletion of *CBF1,* which encodes a master transcriptional regulator of *RP* genes in *C. albicans* but not *S. cerevisiae,* also resulted in a *whi* phenotype. Our analysis also unexpectedly reveals that size regulators may switch between positive and negative functions between the two yeasts. For example, mutation of the conserved transcription factor Dot6 that controls rRNA and *Ribi* expression caused a strong *whi* phenotype in *C. albicans*, in contrast to the large size phenotype conferred in *S. cerevisiae* (27). These results illustrate the evolutionary plasticity of size control mechanisms at the transcriptional level.

In *C. albicans,* the G1/S phase cell cycle machinery remains only partially characterized but nevertheless appears to exhibit disparities compared to *S. cerevisiae.* For instance, despite conservation of SBF and Cln3 function (53, 75), the G1/S repressor Whi5 (15, 18) and the G1/S activator Bck2 (76) appear to have been lost in *C. albicans.* In *S. cerevisiae,* cells lacking *cln3* are viable and able to pass Start due to the redundant role of Bck2 (76), whereas in *C. albicans* Cln3 is essential, presumably due to the absence of a Bck2 equivalent (77). Nrm1 also appears to have replaced Whi5 as it interacts physically with the SBF complex and acts genetically as a repressor of the G1/S transition in *C. albicans* (61). Consistently, we observe that *nrm1* mutant exhibits a reduced cell size as a consequence of accelerated passage through Start. In addition, the promoters of genes that display a peak of expression during the G1/S transition lack the SCB cis-regulatory element recognized by the SBF complex in *S. cerevisiae* and are instead enriched in MCB-like motifs (51).

### Control of Start by the HOG network

Our systematic size screen uncovered a new stress-independent role of the HOG signaling network in coordinating cell growth and division. Hog1 and its metazoan counterparts, the p38 MAPK family, respond to various stresses in fungi (78) and metazoans (79). In contrast to these stress-dependent functions, our data suggests that the basal level activity of the module is required to delay the G1/S transition under non-stressed homeostatic growth conditions. This function of the HOG module appears specific to *C. albicans* as compared to *S. cerevisiae.* Recent work has shown that loss of the metazoan ortholog p38β causes small cell and organism size in *D. melanogaster* (80). In mice, inactivation of the two Hog1 paralogs p38γ and p38δ alters both cell and organ size, including in the heart and the liver (81, 82). Consistently, we have also determined that CRISPR-mediated disruption of p38α or p38β in human cells causes a small size phenotype (data not shown). These observations suggest the role of Hog1 in size control may be more widely conserved and that *C. albicans* may be a suitable yeast model to dissect the mechanisms whereby Hog1 links the growth machinery with cell cycle commitment decision. Here, we show that the entire HOG module is required for cell size control in *C. albicans*, and demonstrate a unique role for the type 2C phosphatases Ptc1 and Ptc2 in size control. In contrast, modulation of the basal activity of Hog1 by the tyrosine phosphatases Ptp2 and Ptp3 in response to a reduction of TOR activity is required for the separate response of hyphal elongation (64). The mechanisms whereby the same MAPK module can specifically respond to stress, nutrient and cell size remains to be resolved.

The signals sensed by the HOG network that couple growth to division also remain to be elucidated. Deletion of the upstream negative regulators of the HOG module, Ypd1 and Sln1, caused an increase in cell size, consistent with the negative regulation of Start by the entire HOG network. Previous studies have suggested that Sln1 histidine phosphotransferase activity is required for cell wall biogenesis in both *S. cerevisiae* and *C. albicans* (83, 84). Interestingly, we also found that disruption of the beta-1,3-glucan synthase subunit Gsc1 also caused a reduced size in *C. albicans.* Based on this result, we speculate that accumulation of cell wall materials, such as glucans, and/or cell wall mechanical proprieties may be sensed through basal activity of the HOG module in order to link growth rate to division. This model is analogous to that postulated in bacteria, whereby the enzymes that synthesize cell wall peptidoglycan help establish cell size control by maintaining cell width (85). In support of this notion, perturbation of the cell wall leads to a G1 phase cell cycle arrest in *S. cerevisiae* via the PKC/Slt2 signalling network (31, 86, 87).

### The HOG network lies at the nexus of growth and cell cycle control

The nature of the linkage between growth to division represents a longstanding general problem in cell biology. The complex genetics of size control, reflected in the hundreds of genes that directly or indirectly affect size, confounds the notion of a simple model of size control (2). Our analysis of Hog1 interactions with the known growth and division machineries nevertheless suggests that the HOG module may directly link growth and division to establish the size threshold at Start. We demonstrate that the HOG module acts genetically upstream of Sfp1 to activate *Ribi* and translation-related genes, and specifically that Hog1 is required for the expression of many genes implicated in ribosome biogenesis and the recruitment of Sfp1 to the relevant promoters. We also demonstrate that Hog1 and its upstream kinase Pbs2 both physically interact with Sfp1, and that Hog1 localizes to many ribosome biogenesis promoters, consistent with a direct regulatory mechanism. These data suggest that basal activity of the HOG module determines ribosome biogenesis and protein synthesis rates. Strikingly, the HOG module also exhibits strong genetic interactions with the SBF transcriptional machinery. The large cell size caused by loss of SBF function is thus epistatic to the small size caused by HOG module mutations. Since Hog1 physically interacts with SBF, the HOG module is ideally positioned to communicate the activity of the growth machinery to the cell cycle machinery. We speculate that under conditions of rapid growth, Hog1 and/or other components of the HOG module may be sequestered away from SBF, thereby delaying the onset of G1/S transcription. In the absence of Hog1 basal activity, this balance is set to a default state, in which SBF is activated prematurely for a given rate of growth. Taken together, these observations suggest a model whereby the HOG module directly links growth to cell cycle commitment (**Figure 8**). The control of SBF by the HOG module appears to operate in parallel to Cln3, Nrm1 and nutrient conditions, suggesting that multiple signals are integrated at the level of SBF, in order to optimize adaptation to different conditions (2). Further analysis of the functional relationships between the HOG module and the numerous other genes that affect size in *C. albicans* should provide further insights into the linkage between growth and division.

### Plasticity of the global size control network and organism fitness

It has been argued that optimization of organism size is a dominant evolutionary force because fitness depends exquisitely on adaptation to a particular size niche (88). The strong link between size and fitness has been elegantly demonstrated through the artificial evolution of *E. coli* strains adapted to different growth rates (3). Comparison of the size phenomes of the opportunistic pathogen *C. albicans* and the saprophytic yeasts *S. cerevisiae* and *S. pombe* reveals many variations in the growth and cell cycle machineries that presumably reflect the different lifestyles of these yeasts. Intriguingly, 30 % of the size regulators identified in our *C. albicans* screen have been previously identified as virulence determinants for this pathogen. This striking correlation suggests that cell size may be an important virulence trait. It is known that other fungal pathogens including *Histoplasma capsulatum, Paracoccidioides brasiliensis, Cryptococcus neoformans* and *Mucor circinelloides* adjust their cell size either to access specific niches in the host or to escape from host immune cells (89). For example, the novel gray cell type recently identified in *C. albicans* is characterized by a small size, a propensity to cause cutaneous infections, and reduced colonization of internal organs (90, 91). Conversely, the host immune system appears to be able to sense *C. albicans* size to modulate the immune response and thereby mitigate tissue damage at the site of infection (92). These lines of evidence underscore the complex relationship between cell size and pathogen fitness. The evident scope and plasticity of the global size control network provides fertile ground for adaptive mechanisms to optimize organism size and fitness.

## Materials and methods

### Strains, mutant collections and growth conditions

*C. albicans* strains were cultured at 30°C in yeast-peptone-dextrose (YPD) medium supplemented with uridine (2 % Bacto peptone, 1 % yeast extract, 2 % w/v dextrose, and 50 mg/ml uridine). Alternative carbon sources (glycerol and ethanol) were used at 2 % w/v. Wt and mutant strains used in this study together with diagnostic PCR primers are listed in **Supplementary file 8**. The kinase (40), the TF (41) and generalist (45) mutant collections used for cell size screens were acquired from the genetic stock center (http://www.fgsc.net). The GRACE (46) collection was purchased from the National Research Council of Canada research center (NRC Royalmount, Montreal). The transcriptional regulator (44) mutant collection was kindly provided by Dr. Dominique Sanglard (University of Lausanne). Growth assay curves were performed in triplicate in 96-well plate format using a Sunrise™ plate-reader (Tecan) at 30°C under constant agitation with OD595 readings taken every 10 min for 24h. TAP and HA tags were introduced into genomic loci as previously described (93). Overexpression constructs were generated with the CIp-Act-cyc plasmid which was linearized with the *StuI* restriction enzyme for integrative transformation (94). The tetracycline repressible mutants *sfp1/pTET-sfp1* and *gpd1/pTET-GPD1* were from the GRACE collection.

### Cell size determination

Cell size distributions were analyzed on a Z2-Coulter Counter (Beckman). *C. albicans* cells were grown overnight in YPD at 30°C, diluted 1000-fold into fresh YPD and grown for 5h at 30°C to a an early log phase density of 5x10^6^ −10^7^ cells/ml. For the GRACE collection, all strains and the wt parental strain CAI-4 were grown overnight in YPD supplemented with the antibiotic doxycycline (40μg/ml) to achieve transcriptional repression. Deletion mutant strains were also grown to early log phase in the presence of doxycycline (40 μg/ml) to allow direct comparisons across the different strain collections. We note that high concentration of doxycycline (100 μg/ml) cause a modest small size phenotype in *C. albicans* but the screen concentration of 40 μg/ml doxycycline did not cause an alteration in cell size. 100 μl of log phase (or 10 μl of stationary phase) culture was diluted in 10 ml of Isoton II electrolyte solution, sonicated three times for 10s and the distribution measured at least 3 times on a Z2-Coulter Counter. Size distributions were normalized to cell counts in each of 256 size bins and size reported as the peak mode value for the distribution. Data analysis and clustering of size distributions were performed using custom R scripts that are available on request.

### Centrifugal elutriation

The critical cell size at Start was determined by plotting budding index as a function of size in synchronous G1 phase fractions obtained using a JE-5.0 elutriation rotor with 40 ml chamber in a J6-Mi centrifuge (Beckman, Fullerton, CA) as described previously (95). *C. albicans* G1 phase cells were released in fresh YPD medium and fractions were harvested at an interval of 10 min to monitor bud index. For the *hog1* mutant strain, additional size fractions were collected to assess transcript levels of the *RNR1*, *PCL2* and *ACT1* as cells progressed through G1 phase at progressively larger sizes.

### Gene expression profiles

Overnight cultures of *hog1* mutant and wt strains were diluted to an OD_595_ of 0.1 in 1 L fresh YPD-uridine media, grown at 30°C to an OD_595_ of 0.8 and separated into size fractions by elutriation at 16°C. A total of 10^8^ G1 phase cells were harvested, released into fresh YPD medium and grown for 15 min prior to harvesting by centrifugation and stored at −80°C. Total RNA was extracted using an RNAeasy purification kit (Qiagen) and glass bead lysis in a Biospec Mini 24 bead-beater. Total RNA was eluted, assessed for integrity on an Agilent 2100 Bioanalyzer prior to cDNA labeling, microarray hybridization and analysis (96). The GSEA Pre-Ranked tool (http://www.broadinstitute.org/gsea/) was used to determine statistical significance of correlations between the transcriptome of the *hog1* mutant with a ranked gene list (97) or GO biological process terms as described by Sellam *et al.* (97). Data were visualized using the Cytoscape (98) and EnrichmentMap plugin (99).

### Promoter localization by ChIP-chip and ChIP-qPCR

ChIP analyses were performed as described using a custom Agilent microarray containing 14400 (8300 intergenic and 6100 intragenic) 60-mer oligonucleotides that covered all intergenic regions, ORFs and different categories of non-coding RNAs (tRNAs, snoRNAs, snRNAs and rRNA (93). A total of 10^7^ G1 phase cells were harvested from log phase cultures by centrifugal elutriation and released into fresh YPD medium for 15 min. Arrays were scanned with a GenePix 4000B Axon scanner, and GenePix Pro software 4.1 was used for quantification of spot intensities and normalization. Hog1 genomic occupancy was determined in duplicate ChIP-chip experiments, which were averaged and thresholded using a cutoff of two standard deviations (SDs) above the mean of log ratios (giving a 2-fold enrichment cutoff). For ChIP analysis of HA-tagged Sfp1, qPCR was performed using an iQ™ 96-well PCR system for 40 amplification cycles and QuantiTect SYBR Green PCR master mix (Qiagen) using 1 ng of captured DNA and total genomic DNA extracted from the whole cell extract. The coding sequence of the *C. albicans ACT1* gene was used as a reference for background in all experiments. Values were calculated as the mean of triplicate experiments.

### Protein immunoprecipitation and immunoblot

Cultures of epitope-tagged strains were grown to OD_595_ of 1.0–1.5 in YPD and either treated or not with rapamycin (0.2 μg/ml) or NaCl (0.5 M) for 30 min. Cells were harvested by centrifugation and lysed by glass beads in IP150 buffer (50 mM Tris-HCl (pH 7.4), 150 mM NaCl, 2 mM MgCl_2_, 0.1 % Nonidet P-40) supplemented with Complete Mini protease inhibitor cocktail tablet (Roche Applied Science) and 1 mM phenylmethylsulfonyl fluoride (PMSF). 1 mg of total protein from clarified lysates was incubated with 50 μl of monoclonal mouse anti-HA (12CA5) antibody (Roche Applied Science), or 20 μl anti-Pbs2 rabbit polyclonal antibody or 20μl anti-Hog1 rabbit polyclonal antibody (Santa Cruz) and captured on 40 μl Protein A-Sepharose beads (GE) at 4*°*C overnight. Beads were washed three times with IP150 buffer, boiled in SDS-PAGE buffer, and resolved by 4–20 % gradient SDS-PAGE. Proteins were transferred onto activated polyvinylidene difluoride (PVDF) membrane and detected by rabbit anti-HA (1:1000) antibody (QED Biosciences) and IRDye680 secondary antibody (LI-COR).

## Acknowledgments

We are grateful to the Fungal Genetics Stock Center (FGSC), Cathrine Bachewich, Ana Traven, Joachim Ernst and Daniel Kornitzer for providing strains. We thank Thierry Bertomeu and Driss Boudeffa for sharing unpublished data. This work was supported by grants from the Natural Sciences and Engineering Research Council of Canada (#06625) to AS, the Canadian Foundation for Innovation to AS and MT, the Canadian Institutes for Health Research (MOP 366608) to MT, the Wellcome Trust (085178) to MT, the National Institutes of Health (R01RR024031) to MT, and from the Ministère de l’enseignement supérieur, de la recherche, de la science et de la technologie du Québec through Génome Québec to MT. JC was supported by a Université Laval Faculty of Medicine and CHUQ foundation PhD scholarships. AS was supported by a start-up award from the Faculty of Medicine, Université Laval and the CHUQ, and by a Fonds de Recherche du Québec-Santé FRQS J1 salary award. MT was supported by a Canada Research Chair Systems and Synthetic Biology.

---

**Figure 1. Figure supplement 1.**
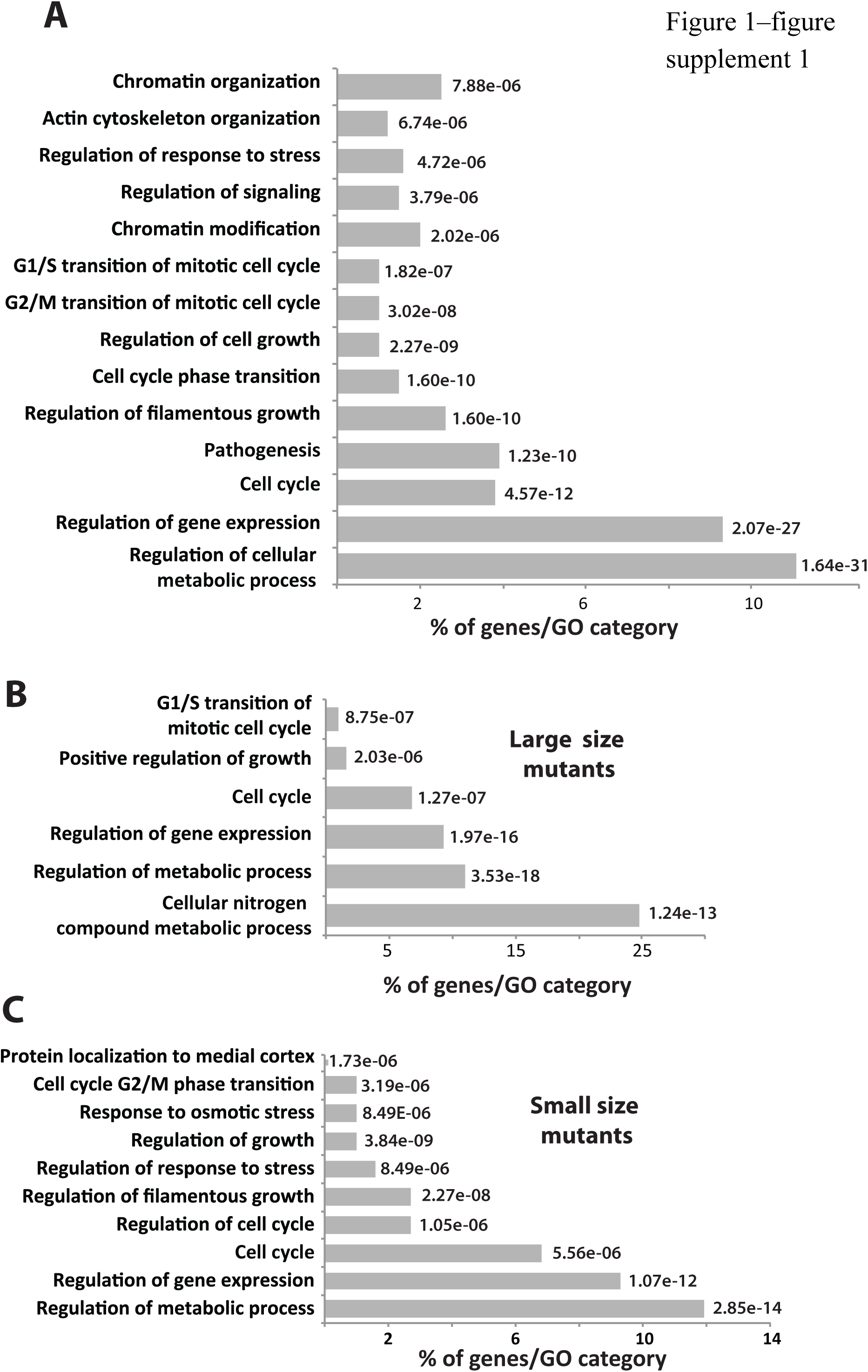
Biological functions of size control genes in *C. albicans.* **(A)** GO biological process term enrichment of all 195 size mutants identified in this study. **(B)** GO term enrichment of small sized mutants. **(C)** GO term enrichment of large sized mutants. p-values were calculated based on a hypergeometric distribution (see http://go.princeton.edu/cG1-bin/GOTermFinder).

**Figure 1. Figure supplement 2.**
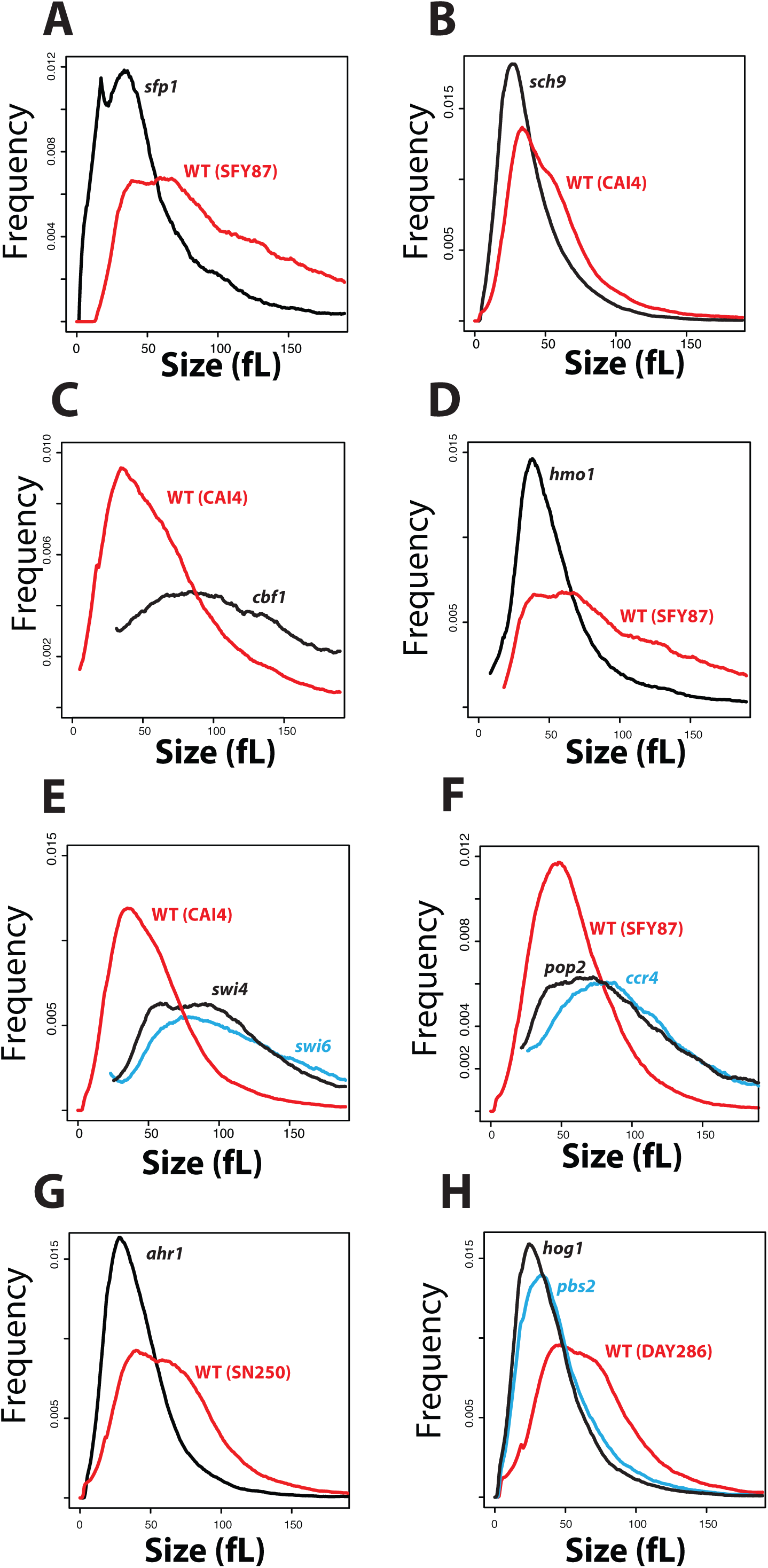
Size distributions of different *C. albicans* size mutants. The indicated mutant strains and congenic wt control strains were grown to early log phase in rich YPD medium and sized on a Z2 coulter channelizer.

**Figure 3. Figure supplement 1.**
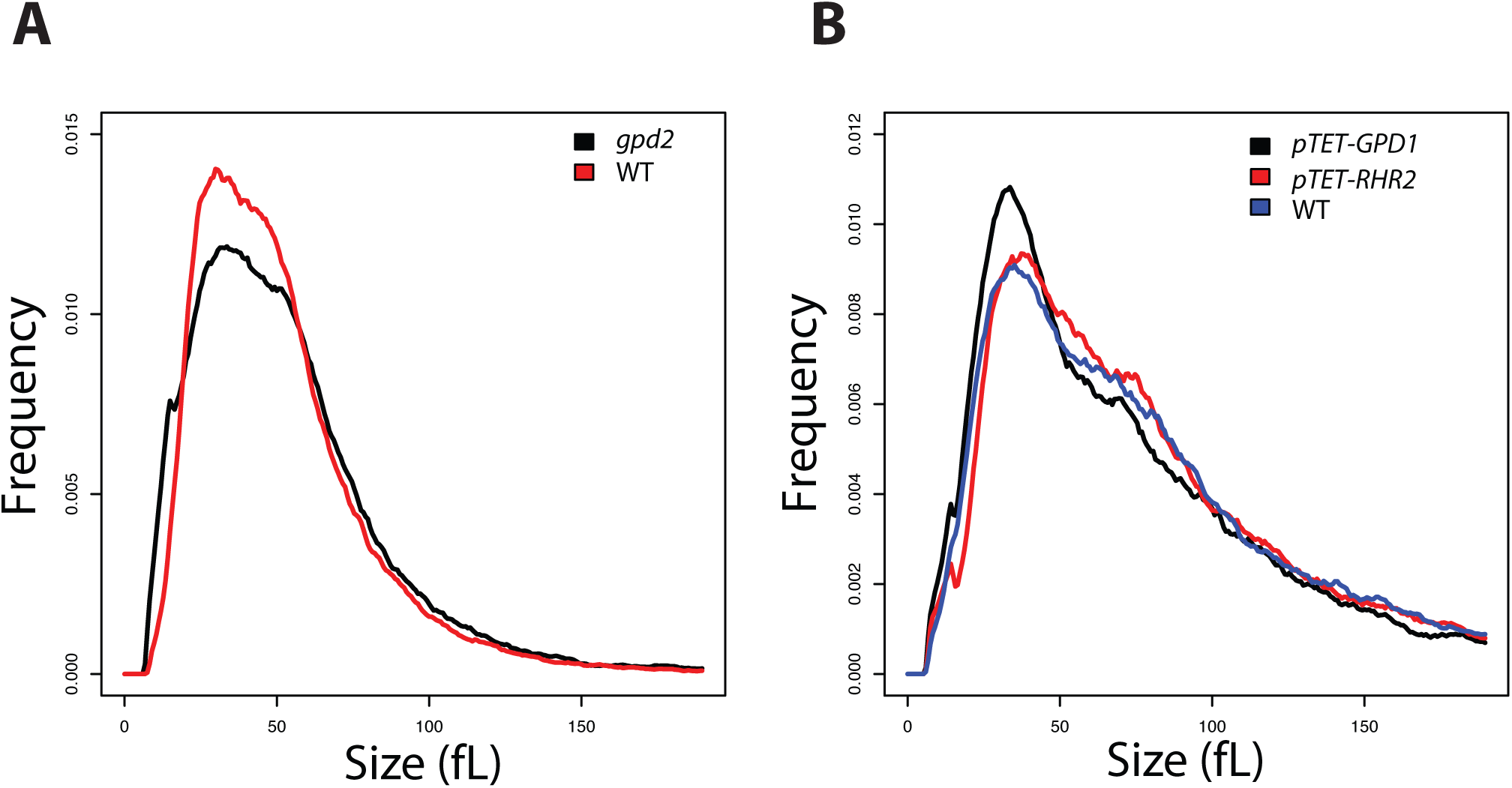
Size distributions of mutants defective in glycerol biosynthesis. **(A)** *gpd2* mutant and wt (SN250) strains were grown to early log phase in rich YPD medium prior to cell size determination. **(B)** The indicated wt (CAI4), *gpd1* and *rhr2* strains were grown to early log phase in rich YPD medium in the presence of doxycycline and sized on a Z2 coulter channelizer.

**Figure 3. Figure supplement 2.**
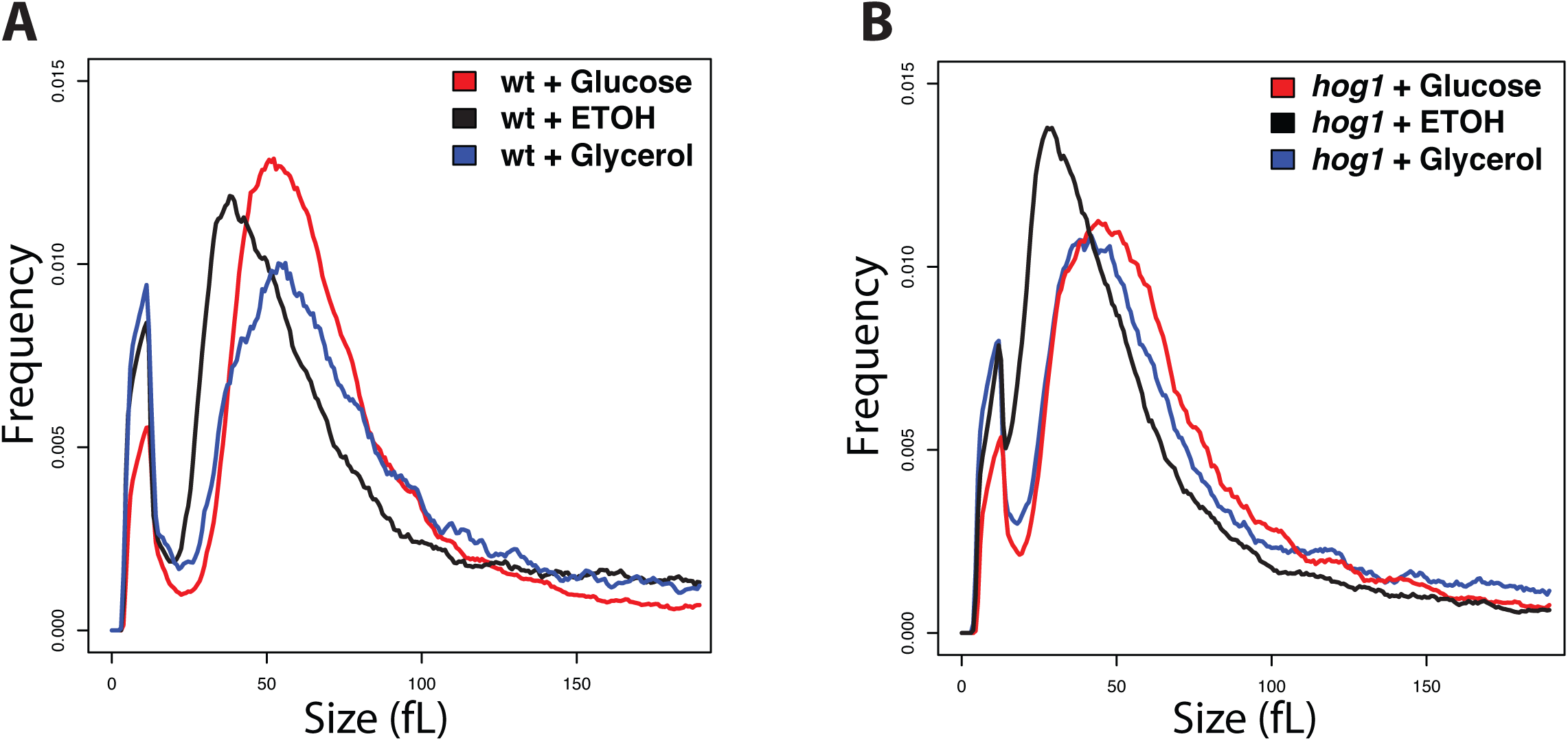
A *hog1* deletion mutant strain adjusts cell size in response to different carbon sources. Size distribution of log-phase cultures of the indicated wt and *hog1* strains (SN250 background) grown in synthetic glucose (red curve), glycerol (blue) and ethanol (black) medium.

**Figure 3. Figure supplement 3.**
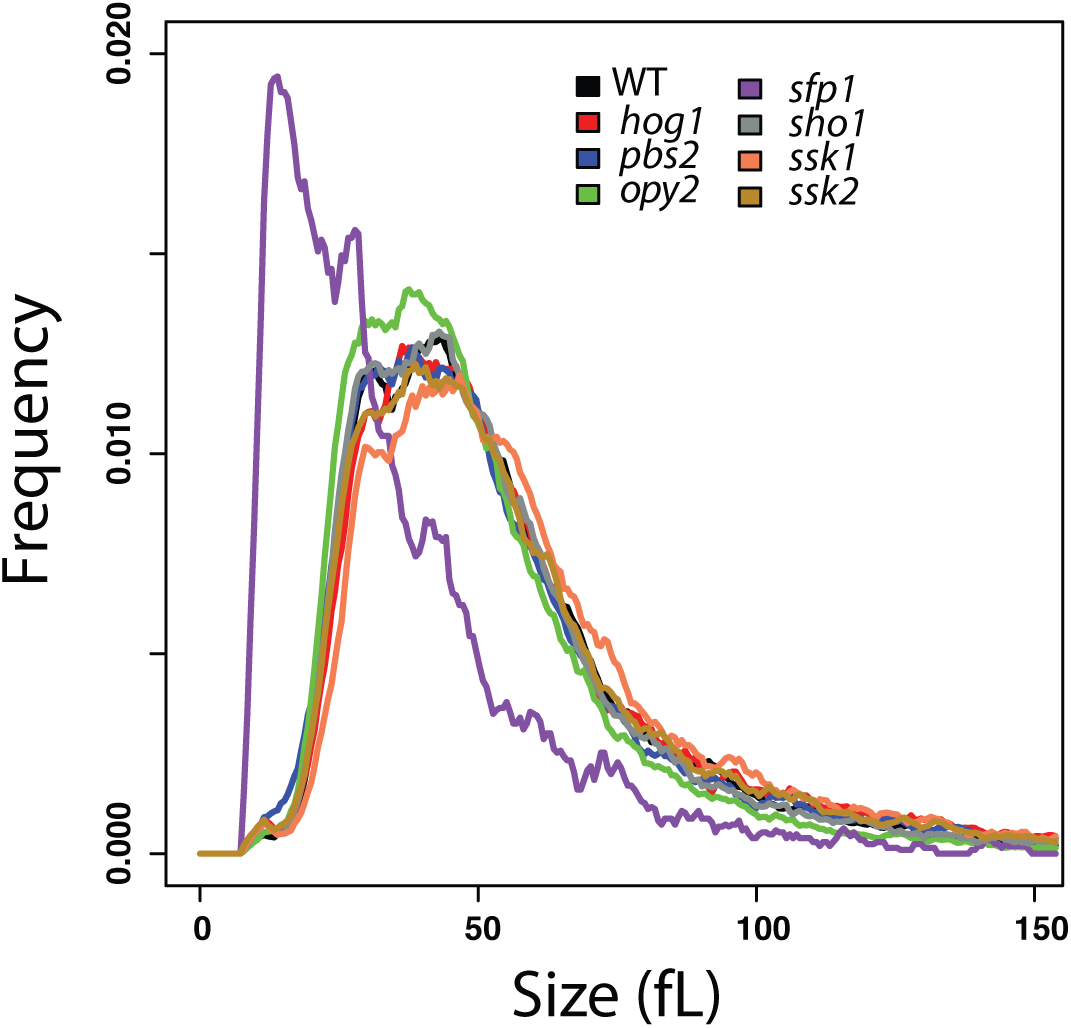
Disruption of central components of *S. cerevisiae* HOG network under non-stressed normo-osmotic conditions. Cultures of the indicated strains were grown to early log phase in rich YPD medium and sized on a Z2 coulter channelizer. Wt (BY4741) and *sfp1Δ* strains were included as controls.

**Figure 7. Figure supplement 1.**
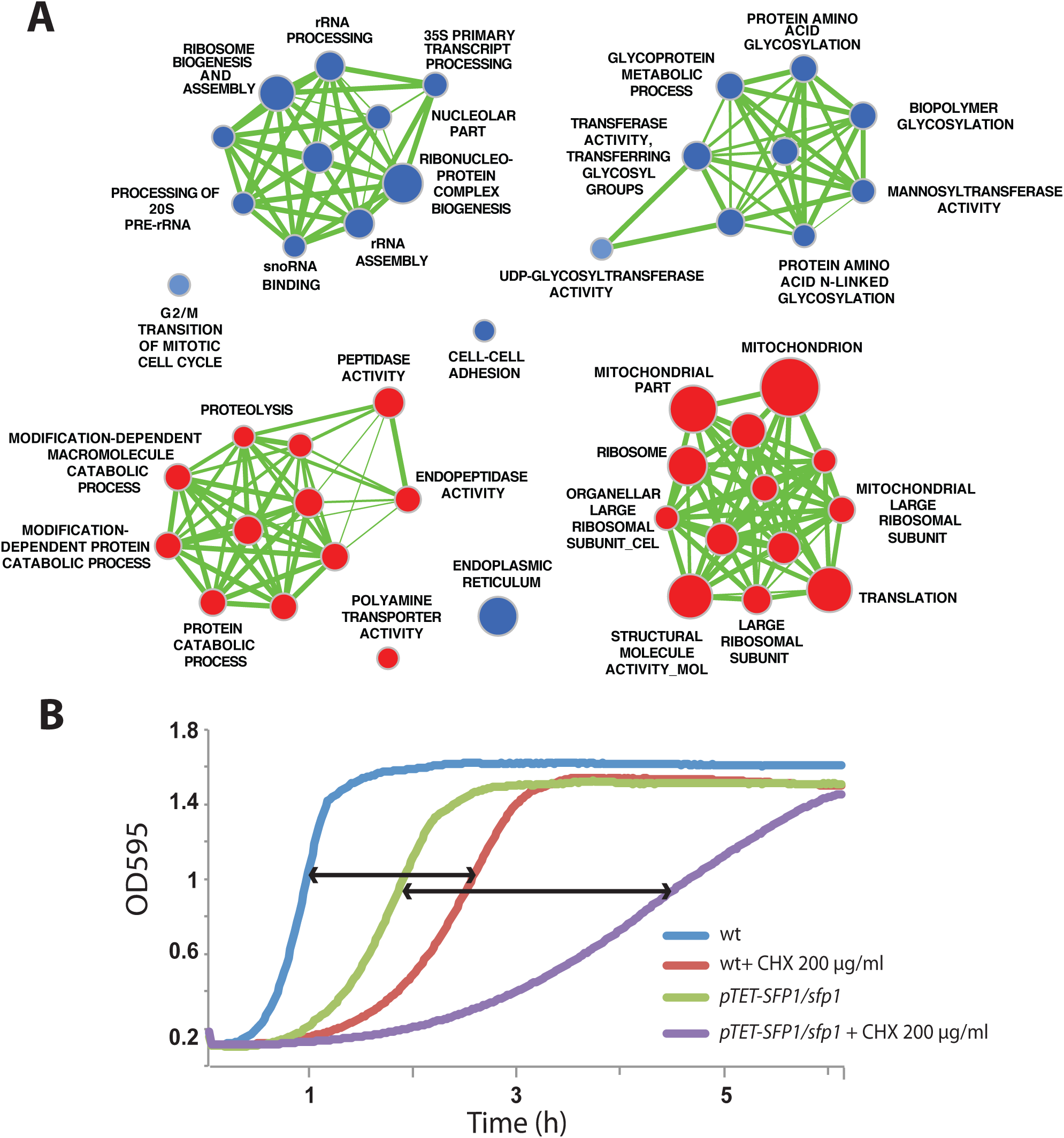
Conservation of Sfp1 function in *C. albicans* as a transcriptional activator of *Ribi* genes. **(A)** Network visualization of transcriptional changes in a tet-*SFP1*/*sfp1* conditional mutant strain. Genes expressed at reduced (blue) or elevated (red) levels after Sfp1 repression were organized into functionally connected networks (green lines) based on Gene Ontology biological process terms. Node size indicates the magnitude of change. Data were visualized using Cytoscape and the Enrichment Map plug-in. **(B)** A pTET-*SFP1*/*sfp1* conditional mutant exhibited increased sensitivity to the protein translation inhibitor cycloheximide (CHX). Cells were grown in YPD at 30°C, and OD_595_ readings were taken every 10 min.

## Additional files

**Supplementary file 1.** Experimental size data of individual mutant strains from the four different *C. albicans* gene deletion collections used in this study. Mean, median and mode size of each strain are indicated.

**Supplementary file 2.** List of 195 size mutants in *C. albicans* that had a greater than 20 % increase or decrease in size compared wt control strains.

**Supplementary file 3.** List of 195 smallest and largest deletion and conditional mutants in *C. albicans* grouped according to GO biological process terms.

**Supplementary file 4.** Doubling-times for small size mutants in *C. albicans.*

**Supplementary file 5.** Gene Set Enrichment Analysis (GSEA) for expression profiles in G1 phase cells determined in a *hog1* strain.

**Supplementary file 6.** Genome-wide promoter occupancy profile of Hog1 in G1 phase cells.

**Supplementary file 7.** Size mutants that exhibited a known virulence defect. Data were extracted from CGD database and reference (100).

**Supplementary file 8**. List of *C. albicans* strains and primers used in this study.

